# Ultrapotent SARS coronavirus-neutralizing single-domain antibodies that bind a conserved membrane proximal epitope of the spike

**DOI:** 10.1101/2023.03.10.531533

**Authors:** Sieglinde De Cae, Inge Van Molle, Loes van Schie, Sophie R. Shoemaker, Julie Deckers, Nincy Debeuf, Sahine Lameire, Wim Nerinckx, Kenny Roose, Daria Fijalkowska, Simon Devos, Anne-Sophie Desmet, Jackeline Cecilia Zavala Marchan, Toon Venneman, Koen Sedeyn, Marlies Ballegeer, Manon Vanheerswynghels, Caroline De Wolf, Hans Demol, Pieter Vanhaverbeke, Gholamreza Hassanzadeh Ghassabeh, Chiara Lonigro, Viki Bockstal, Manuela Rinaldi, Rana Abdelnabi, Johan Neyts, Susan Marqusee, Bart N. Lambrecht, Nico Callewaert, Han Remaut, Xavier Saelens, Bert Schepens

## Abstract

Currently circulating SARS-CoV-2 variants have gained complete or significant resistance to all SARS-CoV-2-neutralizing antibodies that have been used in the clinic. Such antibodies can prevent severe disease in SARS-CoV-2 exposed patients for whom vaccines may not provide optimal protection. Here, we describe single-domain antibodies (VHHs), also known as nanobodies, that can broadly neutralize SARS-CoV-2 with unusually high potency. Structural analysis revealed their binding to a unique, highly conserved, membrane proximal, quaternary epitope in the S2 subunit of the spike. Furthermore, a VHH-human IgG1 Fc fusion, efficiently expressed in Chinese hamster ovary cells as a stable antibody construct, protected hamsters against SARS-CoV-2 replication in a therapeutic setting when administered systemically at low dose. This VHH-based antibody represents a new candidate anti-COVID-19 biologic that targets the Achilles heel of the viral spike.

## Introduction

The spike of SARS-CoV-2 is a major target of neutralizing antibodies. This class I fusion protein consists of a membrane distal S1 and a membrane proximal S2 subunit. The S1 subunit comprises the receptor-binding domain (RBD) and forms the target of the majority of the SARS-CoV-2-neutralizing antibodies available to date (1). The RBD is also immunodominant and tolerates mutations that result in SARS-CoV-2 immune escape (2). As a result of this antigenic evolution, nearly all of the currently emergency use approved monoclonal antibodies fail to neutralize the recently circulating SARS-CoV-2 variants, i.e. BA.4, BA.5, BQ.1.1 and XBB (3, 4). The S2 subunit exerts membrane fusion, a process that involves major conformational changes in the spike, which ultimately results in the formation of a so-called six-helix bundle (6HB) between heptad repeat 1 (HR1) and heptad repeat 2 (HR2) (5). The S2 subunit is more conserved than S1 and, therefore, considered an attractive target for the development of neutralizing antibodies with broad anti-sarbecovirus potential. Several monoclonal antibodies that recognize conserved sites in the S2 subunit such as the stem helix or the fusion peptide of SARS coronaviruses have been described (6–10). In general, however, S2-specific monoclonal antibodies exhibit poor virus neutralizing activity. Single-domain antibodies (VHHs, also known as nanobodies) directed against the SARS-CoV-2 S2 subunit have also been reported, which, even as Fc-fusions, have low SARS-CoV-2-neutralizing activity (11, 12).

Here we report a family of VHHs that can potently neutralize SARS-CoV-2, including past and currently circulating SARS-CoV-2 variants of concern by binding to a highly conserved membrane proximal region in the HR2.

## Results

### Isolation of SARS-CoV-1 and -2 S2 subunit-specific VHHs

To generate a broad repertoire of coronavirus spike-specific VHHs, a llama that had been immunized 4 years earlier with recombinant 2P-stabilized spike proteins derived from Middle-East Respiratory Syndrome coronavirus (MERS-CoV) and SARS-CoV-1 was immunized with recombinant prefusion stabilized SARS-CoV-2-2P spike (13, 14). Subsequently, a VHH-displaying phage library was generated and used for bio-panning. To enrich for VHHs that target the S2 subunit, bio-panning was performed using immobilized SARS-CoV-2-2P spike in the presence of an excess of SARS-CoV-2 RBD in solution. After 3 (R3) or 4 (R4) rounds of bio-panning using either anti-HIS captured SARS-CoV-2-2P spike (R3C and R4C) or directly coated SARS-CoV-2-2P spike (R3DC and R4DC), phagemid clones were selected for small scale VHH production in E. coli. VHH binding to different recombinant coronavirus spike proteins, the SARS-CoV-2 RBD, and the SARS-CoV-2 S2 subunit was determined in ELISA. The VHHs did not bind the SARS-CoV-2 RBD, MERS-CoV or HKU1 spike, but retained binding of SARS-CoV-1 spike (Fig. 1A). Interestingly, the majority of the S2 subunit-binding VHHs could efficiently neutralize vesicular stomatitis virus (VSV) virus particles pseudotyped with SARS-CoV-2 spikes (Fig. 1A). Sequence analysis of the neutralizing VHHs revealed that they belong to a family of 13 unique VHHs with a relatively long CDR3 and that can be further subdivided into 7 subfamilies (fig. S1). At least one VHH from each subfamily was selected, produced in E. coli, purified, and subjected to intact mass spectrometry to confirm their identity. All 10 purified VHHs recognized the full-length spike protein of Wuhan SARS-CoV-2, SARS-CoV-2 BA.1 variant, and SARS-CoV-1 and the S2 subunit of Wuhan SARS-CoV-2 but failed to bind to the SARS-CoV-2 RBD (Fig. 1B). These S2-targeting VHHs also efficiently recognized spike proteins of SARS-CoV-2 D614G, BA.1, BA.2, BA.5, and BQ.1.1 with an intact furin cleavage site expressed on the surface of transfected mammalian cells, whereas MERS-CoV spike was not recognized (Fig. 1C). To evaluate the neutralization potency and breadth of the isolated S2 subunit-targeting VHHs, we performed neutralization assays with VSV pseudotypes displaying spike proteins of SARS-CoV-2 D614G, -BA.2, -BA.5, -XBB, or -BQ.1.1. All VHHs could potently neutralize these SARS-CoV-2 spike pseudotypes with IC_50_ values ranging from 1.1 (R3DC23) to 68.9 ng/ml (R4DC9) (80.0 - 4680 pM) for SARS-CoV-2 both on Vero E6 and VeroE6/TMPRSS2 cells, which allow viral entry at the cell surface (Fig. 1D and E, fig. S2). VHH-R3DC23 was one of the most potently neutralizing S2 binders with an IC_50_ close to 1 ng/ml for SARS-CoV-2 D614G and 2.9-5.8 ng/ml for the tested omicron variants (Fig. 1D and E and fig. S2). The S2-binding VHHs also neutralized replicating VSV SARS-CoV-2 spike pseudotypes with similar potency as the replication-deficient pseudotypes (fig. S3). In line with their binding properties, the isolated VHHs also neutralized VSV pseudotypes carrying the spike protein of SARS-CoV-1 (Fig. 1D and E). Moreover, R3DC23 and R4DC6, which belong to 2 different subfamilies, neutralized authentic SARS-CoV-2 D614G and Omicron BA.1 SARS-CoV-2 viruses with remarkably strong potency (for R3DC23: 4.3 ± 3.5 ng/ml for SARS-CoV-2 D614G and 4.7 ± 2.0 ng/ml for SARS-CoV-2 BA.1, n = 3) (Fig. 1F).

**Figure 1:**
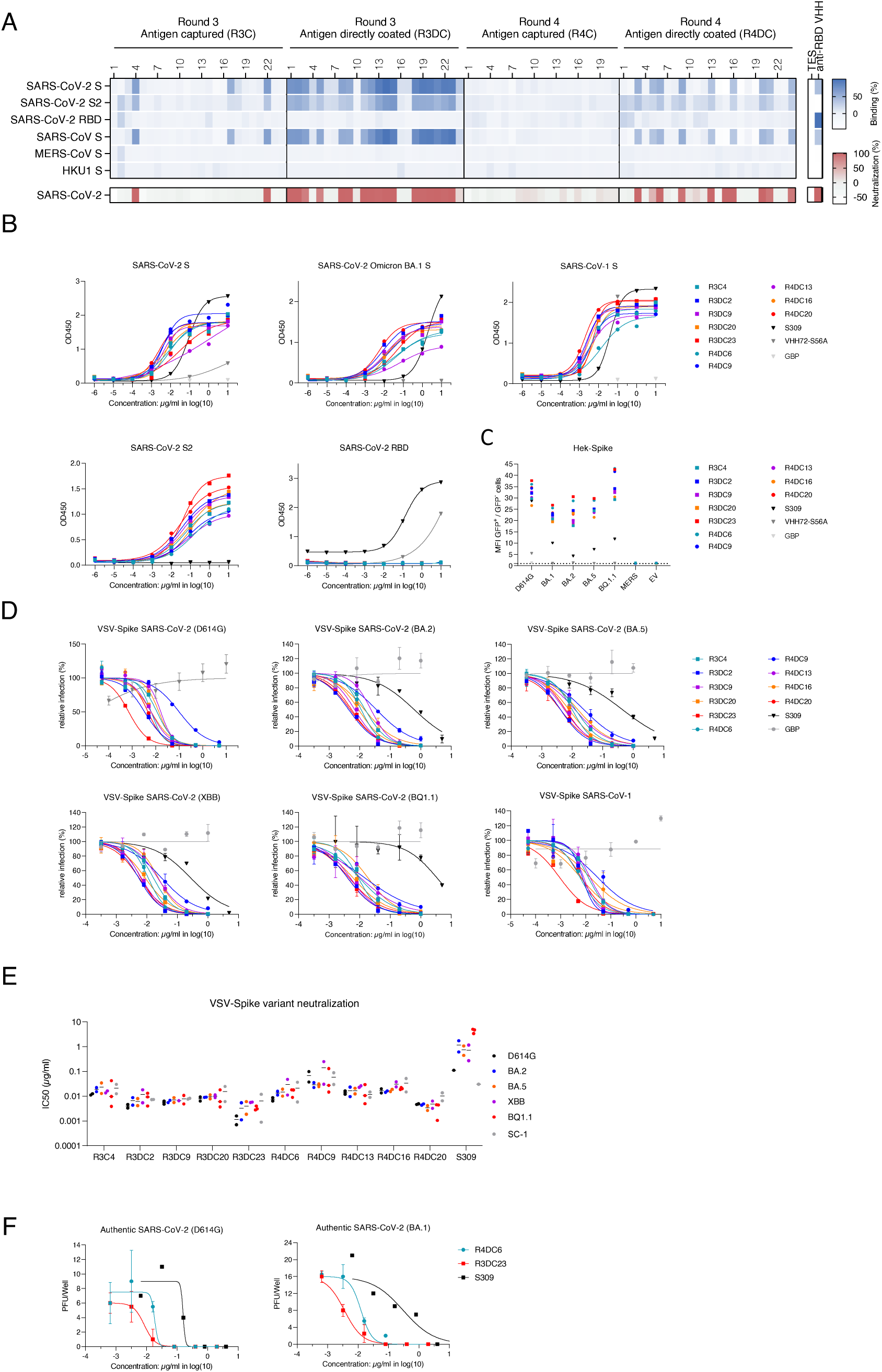
Identification of a VHH family that potently neutralizes SARS-CoV-1 and a broad range of SARS-CoV-2 variants including BQ.1.1 and XBB. **(A)** Screen of E. coli periplasmic extracts (PE) of VHH clones isolated after 3 (R3) or 4 (R4) rounds of bio-panning on SARS-CoV-2 spike protein that was either directly coated (DC) or captured via precoated anti-HIS IgG (C) for binding to the indicated recombinant proteins and for neutralization of VSV particles pseudotyped with SARS-CoV-2 spikes. The blue colored heat map shows for each PE sample (10-fold diluted), the ratio of the ELISA OD_450_signal for the indicated antigen over the ELISA OD_450_ signal of the corresponding PE sample for the control antigen (BSA). PEs prepared from E. coli cells that express an RBD binding VHH, were used as control. Buffer used to prepare the PEs was used as negative control (TES). The red colored heat map shows for each PE sample (100-fold diluted), the level of neutralization of SARS-CoV-2 spike protein pseudotyped VSV. **(B)** Binding of purified VHHs of the R3DC23 family to the spike proteins of SARS-CoV-1, SARS-CoV-2 (with S-6P stabilizing proline mutations) and BA.1, and the S2 subunit (with S-6P stabilizing proline mutations) and the RBD of SARS-CoV-2. The graphs show the OD_450_ signals for the indicated antigens. The GFP binding VHH GBP, RBD specific VHH72-S56A, and S309 were used as controls (14, 54, 55). **(C)** Binding of VHHs to HEK293T cells expressing the spike protein of SARS-CoV-2 D614G, BA.1, BA.2, BA.5, BQ.1.1, or MERS. The SARS-CoV-2 RBD binding VHH72-S56A and S309 were used as positive controls and the GFP binding VHH GBP was used as negative control. The expression of the MERS spike was confirmed by binding of the MERS RBD-specific VHH55 (data not shown) (14). The graph shows the ratio of the MFI of transfected (GFP^+^) cells and the MFI of non-transfected (GFP^-^) cells. **(D and E)** Neutralization of VSV particles pseudotyped with the spike protein of SARS-CoV-2 D614G, BA.2, BA.5, XBB, BQ.1.1 and SARS-CoV-1 by the S2-binding VHHs. The panels in D show representative neutralization assays for each of the tested viruses and the graph in E shows the mean (line) and individual (dots) IC_50_ values calculated from at least 2 independent neutralization assays (E). **(F)** Neutralization of authentic SARS-CoV-2 D614G and BA.1 virus by S2-binding VHHs. The graphs show the mean ± S.D. (N = 2) number of counted plaques for each VHH dilution. S309 was used as positive control.

### S2-binding VHHs prevent spike-mediated membrane fusion

Virus neutralization can be accomplished by preventing receptor-binding and/or, in case of enveloped viruses, membrane fusion. In line with the notion that the S1 subunit is responsible for receptor engagement, S2-targeting VHH R3DC23 did not prevent the binding of purified ACE2-Fc to recombinant SARS-CoV-2 Spike-2P protein (Fig. 2A). Antibodies can, however, also interfere with the hACE2-RBD interaction by prematurely inducing shedding of the S1 subunit (15). Whereas monoclonal antibody CB6 and VHH72_S56A induced S1 shedding, R3C4 and R3DC23 failed to do so (Fig. 2B) (16). In line with this, R3DC23 could not prevent the binding of ACE2-Fc to cells that express SARS-CoV-2 spike whereas VHH72_S56A prevented this interaction (Fig. 2C). These results strongly suggest that our S2-specific VHHs prevent virus entry after attachment. Next, we wondered whether the S2-targeting VHHs could interfere with spike-mediated membrane fusion. Therefore, we tested if R3DC23 could block syncytium formation of VeroE6/TMPRSS2 cells following infection with replication-competent GFP-expressing VSV pseudotyped with SARS-CoV-2 spike. Addition of R3DC23 four hours after infection potently (IC_50_ = 9.2 ± 2.6 ng/ml) prevented syncytium formation whereas moderate inhibition was observed with S309 (IC_50_ = 328.1 ± 110.8 ng/ml) (Fig. 2D). More directly, we also investigated if R3DC23 could prevent cell-cell fusion mediated by mere expression of the spike protein at the cell surface. This was evaluated with Vero E6 cells that had been co-transfected with a SARS-CoV-2 spike and GFP expression vector. Upon spike expression, these cells form large GFP positive syncytia. This spike-mediated syncytium formation was completely prevented by addition of R3DC23 after transfection (Fig. 2E). The impact of R3DC23 on the kinetics of fusion of Vero E6 cells co-expressing spike and GFP was also monitored via time-lapse imaging. In contrast to wells that contain cells that express only GFP, wells that contain cells expressing both GFP and spike proteins displayed a marked increase in the area of GFP-positive cells after 24 hours of transfection, reflecting syncytia formation. This increase was strongly impaired by R3DC23 (Fig. 2F). Based on these observations, we conclude that the S2-targeting VHHs described here can neutralize SARS-CoV-1 and -2 by preventing membrane fusion.

**Figure 2:**
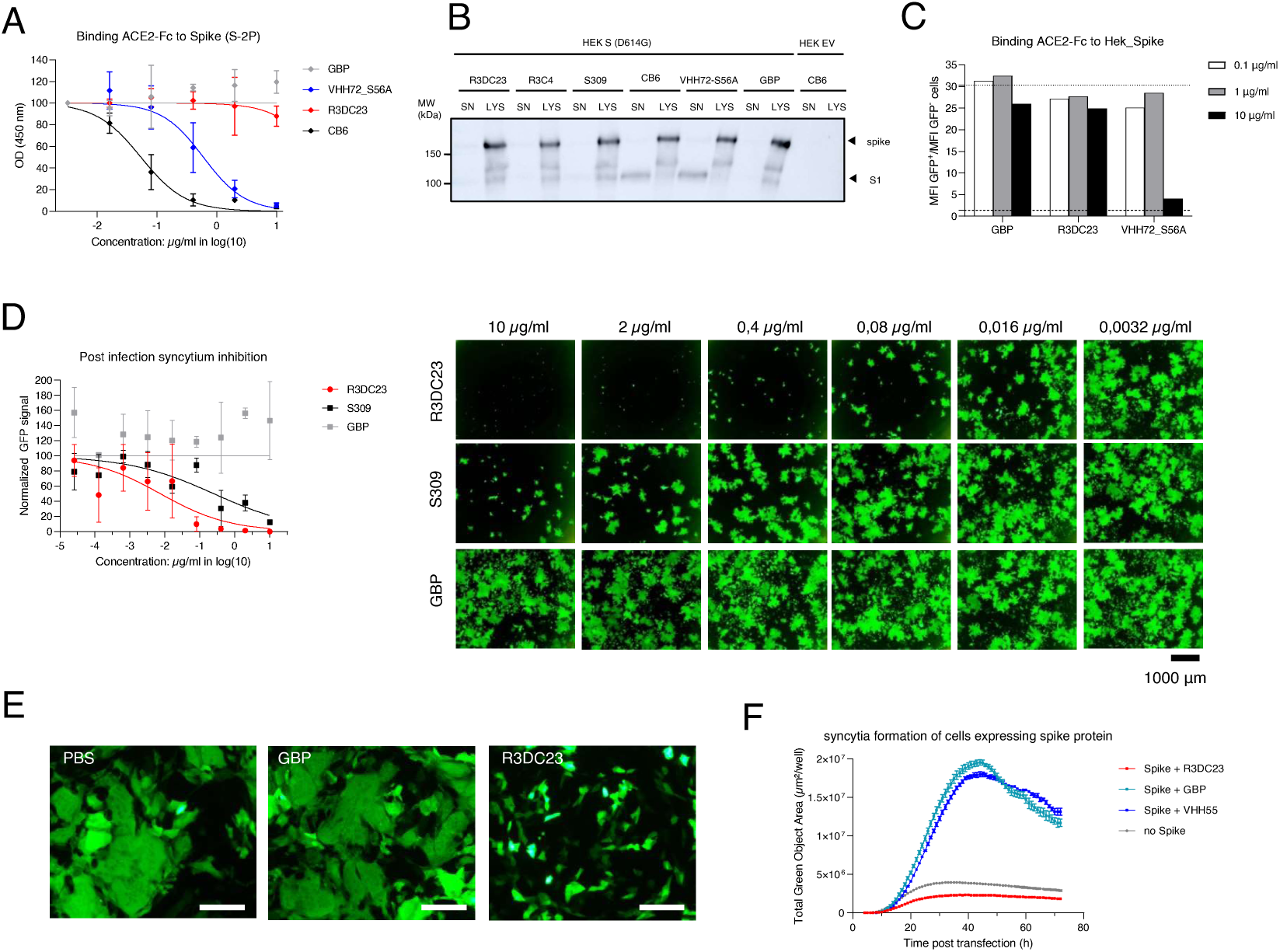
S2-targeting VHHs inhibit spike-mediated membrane fusion. **(A)** S2-targeting VHHs do not interfere with the binding of the spike protein with ACE2. The graph shows the mean ± SD (N = 2) OD_450_ signal of an ELISA in which the binding of human ACE2-muFc to coated recombinant 6P stabilized spike proteins containing an inactivated furin cleavage site was tested in the presence of a dilution series of R3DC23. The GFP binding VHH (GBP) was used as negative control and VHH72-S56A and CB6 as positive controls. **(B)** R3DC23 does not induce shedding of the S1 subunit. The panel shows anti-S1 Western blot analysis of the growth medium (SN) and cell lysates (LYS) of HEK293T cells expressing the SARS-CoV-2 spike protein with D614G substitution (HEK S (D614G)) or not (HEK EV) incubated for 30 minutes with the indicated VHH constructs. The CB6 and S309 antibodies, know to respectively evoke and not to evoke S1 shedding were used as controls. Triangles at the right indicate cell-associated uncleaved spike and the S1 spike subunit generated after furin-mediated cleavage of the spike protein. **(C)** R3DC23 does not interfere with the binding of human ACE2-muFc to cells expressing the spike protein with intact furin cleavage site. The graph shows the ratio of MFI (detection of cell-bound ACE2-muFc) of GFP^+^ cells over that of GFP^-^ cells in the presence of R3DC23. GBP was used as negative control and VHH72-S56A that induces S1 shedding and competes with human ACE2 for the binding to the RBD was used as positive control. The dashed line represents the binding of ACE2-muFc to cells that do not express spike. The dotted line represents the binding of ACE2-muFc to spike expressing cells in the absence of antibody **(D)** R3DC23 prevents syncytia formation by infected cells. The graph shows the mean ± SD (N = 2) GFP fluorescence of wells of Vero E6 cells treated with dilutions series of R3DC23, GBP or S309 4 hours after infection with GFP expressing replication-competent VSV pseudotyped with SARS-CoV-2 spikes. The images on the right show the GFP expression of the indicated samples at 40 hours post-infection. **(E)** R3DC23 potently prevents syncytia formation by cells transfected with a spike protein expression plasmid. The images show the GFP expression of Vero E6 cells that were treated with PBS, GBP or R3DC23 4 hours after co-transfection of a GFP and a spike expression vector. The scale bar represents 250 μm. **(F)** Quantification of syncytia formation by spike expressing cells in the presence of R3DC23, GBP or PBS with live cell imaging. The graph shows the mean ± SD (N = 3) GFP-positive area of wells treated with the indicated VHHs 2 hours after co-transfection of an GFP and a spike expression vector or of only a GFP expression vector (no Spike).

### S2-binding VHHs bind to the membrane proximal region of heptad repeat 2

We performed escape virus selection experiments to better understand the mechanism by which R3DC23 inhibits spike-mediated membrane fusion and, also, as a first approach to identify its likely binding region in the S2 subunit. R3DC23 can potently inhibit syncytia formation of infected cells (Fig. 2D). Therefore, escape virus selection was performed by infecting VeroE6/TMPRSS2 cells with replication-competent GFP-expressing VSV pseudotyped with SARS-CoV-2 spikes in the presence of R3DC23 or an irrelevant control VHH, added 2 hours after infection. Rapid syncytium formation and spread of the infection to neighboring cells was observed in wells to which the control VHH had been added, whereas these events were blocked or strongly delayed in wells with R3DC23. Nine escape viral clones derived from wells with delayed syncytia in the presence of R3DC23 were isolated by limiting dilution, and their spike coding sequence was determined. Each of these viral clones had acquired a single amino acid substitution at one out of four positions: N1192D (1 clone), L1197P (2 clones), L1200P (1 clone), Q1201R (4 clones), and Q1201K (1 clone), all located within the membrane proximal region of HR2 (Fig. 3A and B). Escape viruses with mutations L1197P, L1200P, or Q1201R were completely resistant to R3DC23 neutralization whereas the N1192D mutant virus remained sensitive to neutralization by R3DC23, especially on Vero E6 cells (Fig. 3C and D). In line with this, R3DC23 could still bind to cell surface expressed spike with the N1192D substitution whereas binding to spike mutants with any of the other escape selection mutations was lost (Fig. 3E).

**Figure 3:**
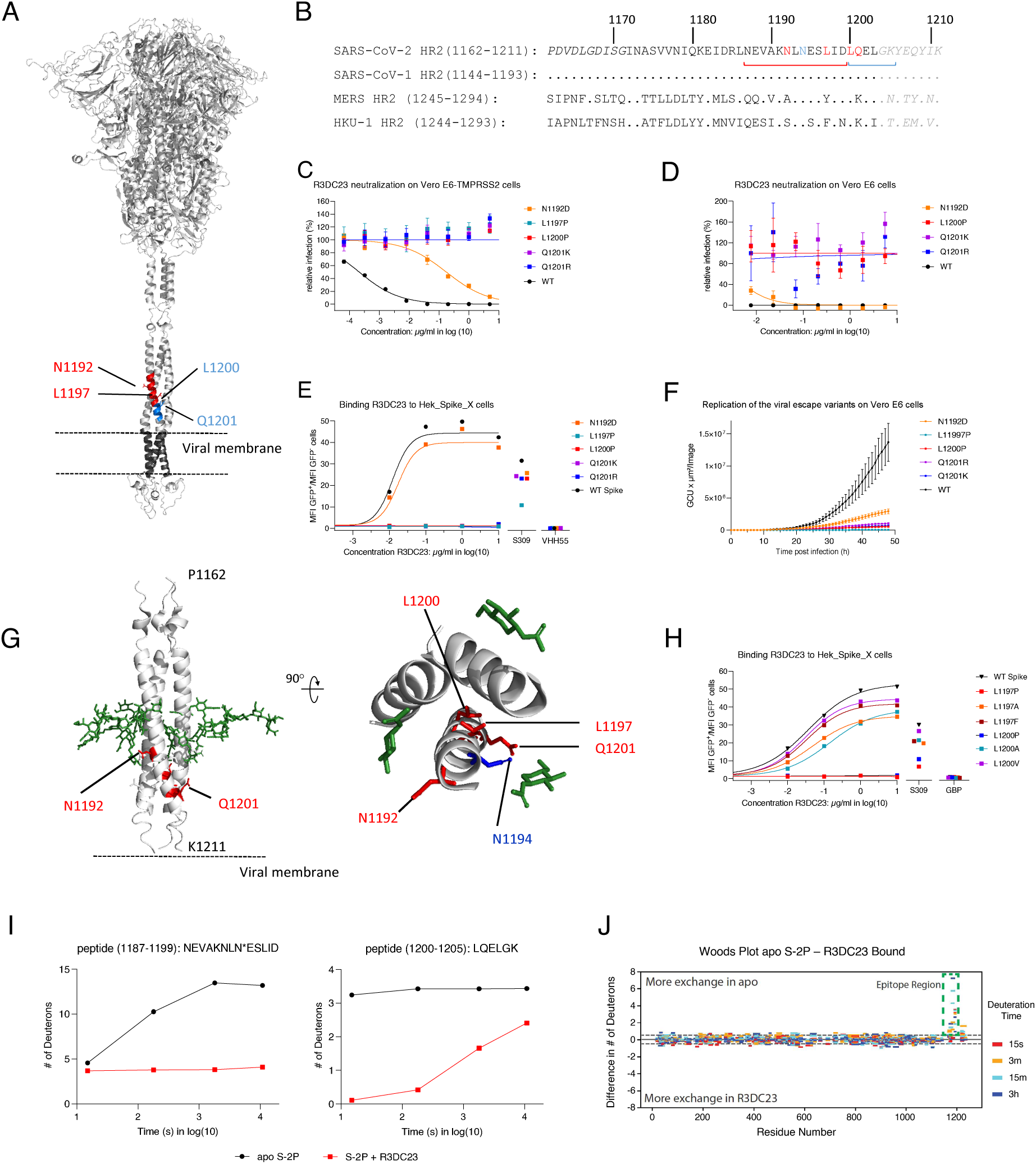
VHH R3DC23 binds the spike protein at a membrane proximal site in the HR2 region. **(A and B)** Replication of VSV virus pseudotyped with Wuhan SARS-CoV-2 spikes on VeroE6/TMPRSS2 cells in the presence of R3DC23 selects viruses with substitutions at 4 positions at the membrane proximal region of the HR2. Panel A shows the complete spike trimer (6XVV_1_1_1) with the amino acids at which substitutions were observed in the escape viruses indicated as red and blue sticks on one spike protomer. The region corresponding to the two peptides in the spike that were protected from HDX by R3DC23 are indicated in red and blue on one spike protomer. Panel B shows the HR2 amino acid sequence of SARS-CoV-2, SARS-CoV-1, MERS and HKU1. The dots represent amino acids that are identical to SARS-CoV-2 HR2. The positions at which substitutions were observed in the escape variants are indicated in red and the N-glycosylation site is indicated in blue. The sequence in normal font specifies the part of the HR2 that contains the HxxHxxx heptad repeat motif (H: a hydrophobic amino acid). The part in grey italic is the flexible linker between the HR2 and transmembrane region. The sequence indicated by the red and blue braces are the two spike peptides that were protected by R3DC23 from HDX. **(C and D)** Replication of viral escape variants N1192D, L1197P, L1200P, Q1201R and Q1201K on VeroE6/TMPRSS2 and Vero E6 cells in the presence of R3DC23. The graphs show the mean ± SEM (N = 4) level of GFP normalized by the GFP fluorescence of mock infected cells and infected cells in the absence of R3DC23. **(E)** Binding of R3DC23 to cells expressing the Wuhan and N1192D, L1197P, L1200P, Q1201R, or Q1201K spike variants. The graph shows the ratio of the MFI of transfected (GFP^+^) cells and the MFI of non transfected (GFP^-^) cells stained with the indicated concentrations of R3DC23, with 10 μg/ml of VHH55 or 1 μg/ml of S309. **(F)** Kinetics of viral replication of replication-competent VSV pseudotyped with SARS-CoV-2 Wuhan (parental) or the selected escape variants as measured by life cell imaging of infected Vero E6 cells. The graph shows the mean ± SEM (N = 5) GFP^+^ area per well of infected cells at the indicated time points post-infection. **(G)** The HR2 coiled-coil structure (PDB:2FXP) as determined by NMR on which the positions at which a substitution was observed in the escape viruses are indicated in red. The glycans conjugated at N1194 as modeled in 6XVV_1_1_1 are indicated in green. The dashed line represents the viral membrane. The right panel represents a top view of part of the HR2 (K1191-K1205) coiled-coil with the N1192, L1197, L1200, and Q1201 indicated in red and N1194 in blue on one protomer. The first GlcNAc of the N-glycans conjugated to N1194 as modelled in 6XVV_1_1_1 are indicated in green (56). **(H)** Binding of R3DC23 to cells expressing the indicated SARS-CoV-2 spike protein variants. The graph shows the ratio of the MFI of transfected (GFP^+^) cells and the MFI of non-transfected (GFP^-^) cells stained with the indicated concentrations of R3DC23, with 10 μg/ml GBP, or 1 μg/ml S309. **(I and J)** Identification of the R3DC23 binding region on recombinant spike protein by HDX-MS. The panels in I show the HDX-MS uptake plots of the two peptides with high degrees of protection from deuteration upon the binding of R3DC23. The asterisk indicates the glycosylated residue. The glycosylated peptide has taken up more deuterium than the number of backbone exchangeable sites (11 sites) because the glycan can uptake and retain deuterium at amide sites similarly to the backbone as noted by Guttman, Scian and Lee (57). (J) Woods plot with each peptide (indicated by the residues numbering in the x-axis) for the indicated time points the difference in the number of deuterons acquired between apo spikes and R3DC23 bound S-2P spikes.

The amino acids in the escape viruses that abolish R3DC23 binding are confined to the part of the spike that sits between the viral membrane and a large tetra-antennary N-glycan at position N1194 (17). The complete SARS-CoV-2 HR2 sequence is identical to that of SARS-CoV-1 but differs in MERS and HKU1, explaining the lack of binding to the spikes of the latter viruses (Fig. 3B). In the SARS-CoV-2 spike prefusion conformation the membrane proximal part of HR2 assembles into a coiled-coil of 3 parallel alpha helices that, upon transition to the postfusion conformation, initially disassemble and then bring the viral and host cell membrane in proximity by zippering up with the HR1 coiled-coil to form a stable 6HB (18). This crucial function of the membrane proximal HR2 region in the fusion process may explain why the VSV pseudotype R3DC23 escape viruses were severely attenuated (Fig. 3F). According to the available NMR structure of SARS-CoV-1 HR2, Q1201 is oriented outwards but both L1197 and L1200 are oriented towards the center of the coiled-coil (Fig. 3G). This suggests that R3DC23 either binds an open conformation of the HR2 or that the leucines at positions 1197 and 1200 do not make direct contact with R3DC23 but that substitutions at these positions to prolines have a significant impact on the folding of the HR2 coiled-coil. To address this, we evaluated R3DC23 binding to spike muteins with either a L1197F, L1197A, L1200V, or L1200A substitution instead of a proline, which is considered a helix breaker (19). R3DC23 could bind to spike with L1197F, L1197A, L1200V, or L1200A substitution, indicating that an intact HR2 coiled-coil tertiary structure is essential for its binding (Fig. 3H).

As another approach to identify the R3DC23 binding region, hydrogen-deuterium exchange monitored by mass spectrometry (HDX-MS) was conducted on recombinant SARS-CoV-2 spike in the presence and absence of R3DC23. Two adjacent peptides from residues 1187 through 1205 were found to be highly protected in the presence of R3DC23 (Fig. 3I). Importantly, all 4 positions at which viral escape variants acquired mutations locate within those two peptides. HDX was not altered by R3DC23 binding in any other regions of the spike (Fig. 3J). These data indicate that R3DC23 binds to the membrane proximal region of HR2.

### R3DC23 recognizes a quaternary epitope in HR2

To get detailed insight in the interactions between R3DC23 and its target we resolved the crystal structure of R3DC23 in complex with a peptide spanning the complete HR2 (H1159-K1211). The crystal asymmetric unit shows an HR2 coiled-coil trimer in complex with three R3DC23 molecules, each binding the interface between two HR2 peptides (Fig. 4). The R3DC23 binding site spans residues N1192 to Y1206, encompassing the C-terminal region of HR2 (Fig. 4A-C). R3DC23 binds two adjacent HR2 helices, encompassing a 407 Å^2^ buried surface area, 8 H-bonds, 2 salt bridges and a calculated solvation free energy gain Δ^i^G for complex formation (i.e. hydrophobic contribution to binding) of -4.7 kcal/mol for helix (i), and a 461 Å^2^ buried surface area, 6 H-bonds, 1 salt bridge and a calculated Δ^i^G of -2.4 kcal/mol for helix (ii) (Fig. 4C, Supplementary Fig. 4). The VHH CDR3 forms the dominant contact surface in the complex, where N100a and Y100b form an extensive H-bond network with N1194, Q1201 and E1202 in helix (ii) and S1196 in helix (i), Y96 goes in H-bond contact with helix (ii) main chain carbonyls, S98 goes in H-bond interaction with E1195 in helix (i), and V97 binds a hydrophobic patch formed by L1200 and L1203 in helix (i). The VHH CDR1 and CDR2 are involved, respectively, in hydrophobic interactions and two salt bridges (R52 – D1199) with helix (i) (Fig. 4C, Supplementary Fig. 4). The identification of Q1201 as a R3DC23 escape mutant agrees with its central part in the H-bond network with CDR3. L1197 and L1200, however, are not in direct contact with R3DC23, suggesting that substitution by proline at these positions contribute indirectly to escape, likely as a result of a destabilization and/or conformational adjustment of the adjoined binding epitope across the HR2 coiled-coil interface. The conformation of the HR2 trimer in complex with R3DC23 closely matches that seen in the pre-fusion form of the S protein, where it corresponds to an approximately 3 nm high region between the membrane and the HR2 and linker regions covered by N-glycosylation (Fig. 4A). In the postfusion form, HR2 helices rearrange to bind the surface of a HR1 coiled-coil, thereby breaking the pairwise HR2 contacts and resulting in partial unfolding of the HR2 C-terminal region (Fig. 4A, D). This rearrangement disrupts the R3DC23 binding site (Fig. 4D). Conversely, the interfacial binding of R3DC23 to the HR2 coiled-coil likely stabilizes the pre-fusion S protein.

**Figure 4:**
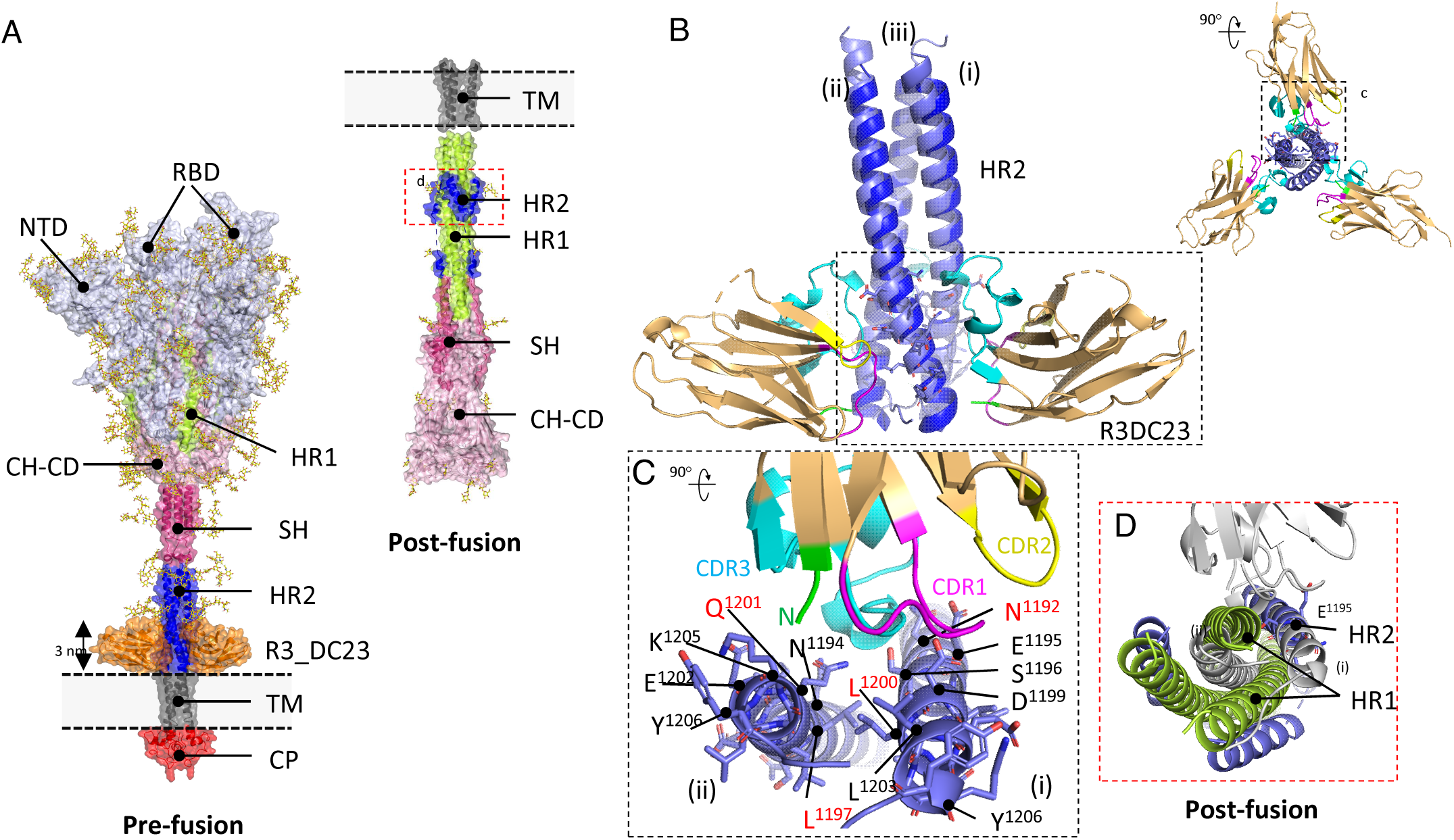
X-ray structure of the R3DC23 – HR2 complex. **(A)** left: model of the full length prefusion S protein (6VSB_1_1_2; (56)) in superimposition with the R3DC23 – HR2 complex, all shown in molecular surface representation, with N-glycans in stick representation. R3DC23 VHHs are colored orange, labelled S protein regions: cytoplasmic domain (red; CP), transmembrane domain (grey; TM), heptad repeat 2 (blue; HR2), S2 stem helix (deep pink; SH), heptad repeat 1 (lemon, HR1), central helix and connector domain (pink; CH-CD). The S1 regions encompassing the N-terminal domain (NTD) and receptor binding domain (RBD). Right: model of the proteolytically processed postfusion S protein (7E9T; (58)), color coded as prefusion spike. **(B)** Side and axial view (inset) of the R3DC23 – HR2 complex (sand and blue, resp.) superimposed with prefusion HR2 coiled coil (light blue). In R3DC23, CDR1, 2, and 3 are colored magenta, yellow, and cyan, resp. The HR2-binding epitope spanning D1192 – Y1206 is shown in stick representation. **(C)** close-up of boxed region, encompassing a single VHH and two HR2 copies (i and ii) forming the adjoined binding epitope. Escape mutant positions are labeled in red. **(D)** axial view of HR1-HR2 (lemon and blue) region of postfusion S protein, superimposed with R3DC23 in the HR2 complex (grey).

### Humanized R3DC23 fused to human IgG1-Fc broadly neutralizes SARS-CoV-2 variants and protects hamsters

With clinical development in mind, we humanized the frame work regions of R3DC23 and two related VHHs (R3C4 and R4DC20) and replaced the N-terminal glutamine by an aspartate residue (16). In addition, these S2-binding VHHs were genetically fused to a human IgG1-Fc_YTE, which creates bivalency and is a well-established approach to extend the in vivo half-life in circulation of a biologic. The resulting humanized constructs, respectively named huR3DC23-Fc, huR3C4-Fc and huR4DC20-Fc were produced in mammalian cells and compared with their monovalent counterpart for neutralization of pseudotyped VSV displaying the spike protein of SARS-CoV-2 D614G and BA.5. For each of these VHHs the humanized Fc fusions neutralized SARS-CoV-2 D614G and BA.5 more efficiently than their monovalent formats with huR3DC23-Fc being the most potent. Moreover, humanization of R3DC23 did not affect the neutralizing activity of huR3DC23-Fc (Fig. 5B). An important parameter for the development of biologicals is their solubility. Therefore, we investigated the hydrophobicity of the VHH-Fc fusions by hydrophobic interaction chromatography. Of the three humanized VHH-Fc fusion constructs the retention times on hydrophobic interaction chromatography was shortest for huR3DC23-Fc, below that of its non-humanized counterpart and well below that of clinically validated VHH-Fc XVR011 (Fig. 5C). These data indicate that, based on solubility and neutralizing activity huR3DC23-Fc, is favorable for further development. Similar as for SARS-CoV-2 D614G and BA.5, huR3DC23-Fc could also potently neutralize VSV pseudotypes displaying the spike protein of SARS-CoV-2 BA.1, BA.2, BA.2.75, BA.4.6, BQ.1.1, and XBB with IC_50_values close to or below 1 ng/ml (Fig. 5D and E). Moreover, huR3DC23-Fc could also potently neutralize authentic SARS-CoV-2 D614G and BA.1 (Fig. 5F).

**Figure 5:**
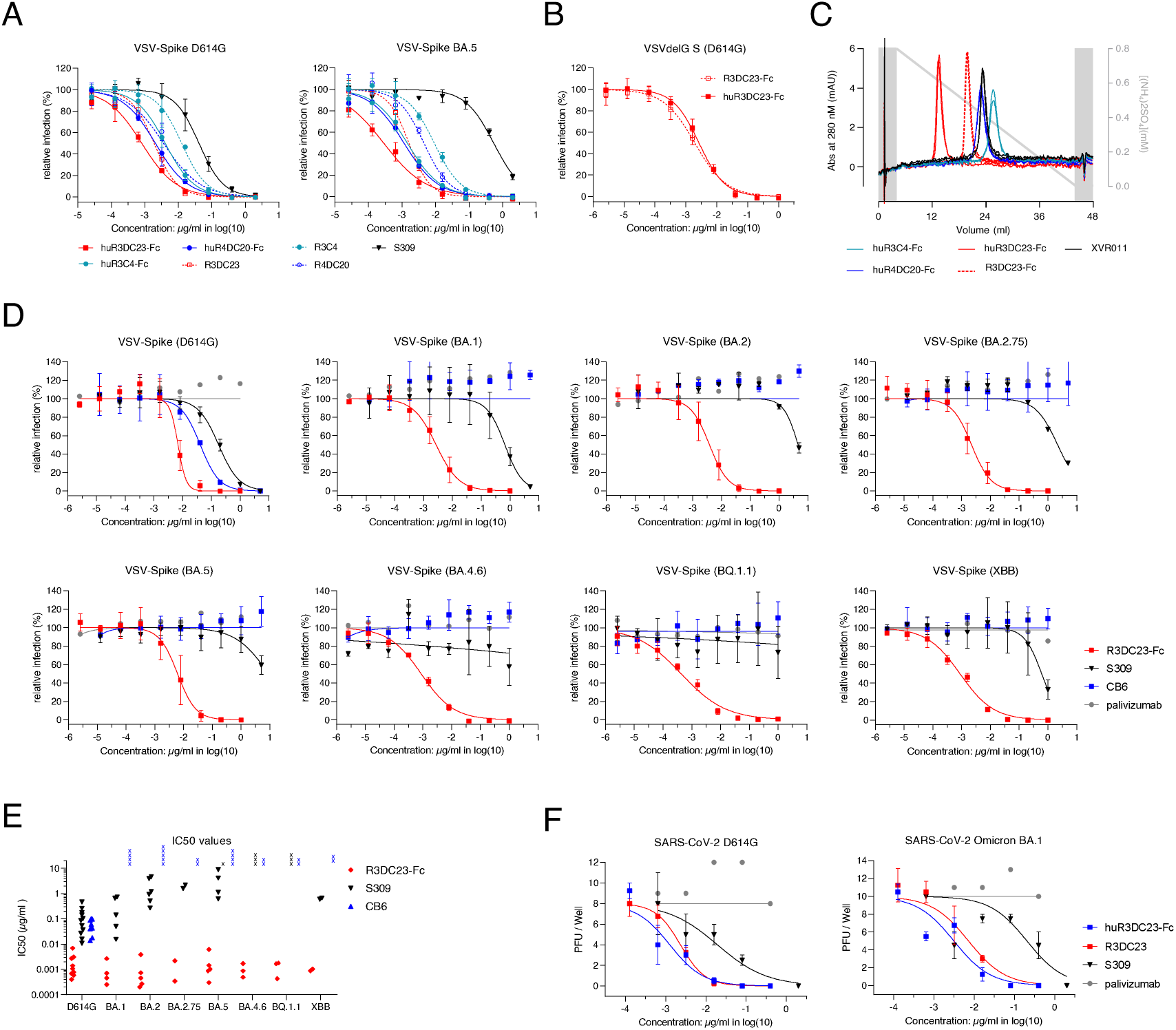
R3DC23-Fc fusions potently neutralize a broad range of SARS-CoV-2 variants. **(A)** Neutralization of SARS-CoV-2 D614G and BA.5 spike VSV pseudotypes by monovalent R3DC23, R3C4, and R4DC20 and their corresponding humanized counterparts fused to human IgG1 Fc containing the half-life extending YTE mutation (huR3DC23-Fc, huR3C4-Fc, and huR4DC20-Fc). The symbols represent the mean ± SD (N = 3) relative infection as measured by GFP fluorescence of infected cells. **(B)** Neutralization of SARS-CoV-2 D614G spike VSV pseudotypes by Fc fusions of humanized (huR3DC23-Fc) and non-humanized (R3DC23-Fc) R3DC23. **(C)** Analytical hydrophobic interaction chromatography of R3DC23-Fc, huR3DC23-Fc, huR3C4-Fc, and huR4DC20-Fc as compared to that of clinically validated VHH-Fc XVR011. Apparent hydrophobicity was assessed on ProPac HIC-10 HPLC over an (NH_4_)_2_SO_4_ elution gradient (short retention times indicate low apparent hydrophobicity). The panel shows duplicate curves for each indicated VHH-Fc construct. **(D)** Neutralization of VSV pseudotyped with spikes of the indicated SARS-CoV variants by huR3DC23-Fc, CB6 and S309. The representative graphs show the mean ± SD (N = 3 for huR3DC23, N = 2 for CB6 and S309) relative infection as measured by GFP fluorescence of infected cells. **(E)** The graph shows the median (line) and individual (diamond) IC_50_ values calculated from at least 2 independent neutralization assays using VSV pseudotyped with the spike protein of the indicated SARS-CoV-2 variants. **(F)** Neutralization of authentic SARS-CoV-2 D614G and BA.1 virus. The graphs show the mean ± SEM (N = 4 for R3DC23 and huR3DC23-Fc and N = 2 for S309) number of counted plaques for each VHH dilution.

LS (M428L/N434S) substitutions in Fc also increase the half-life of engineered antibodies by increasing their affinity for FcRn at low pH, reminiscent of the endosomal compartment. Therefore, we also fused humanized R3DC23 to an Fc fragment with the LS substitutions to obtain huR3DC23-Fc_LS. huR3DC23-Fc_LS displayed low apparent hydrophobicity and eluted in a single peak from SEC in between 44 and 158 kDa molar weight markers, consistent with a fully assembled VHH-Fc (Fig. 6A and -B). To confirm that also in VHH-Fc fusions the LS mutation is associated with a high affinity of FcRn, the affinity of huR3DC23-Fc-LS for human FcRn was tested at pH6. At 3.5 - 4.5 nM (steady state affinity and 1:1 binding model, respectively), the affinity of huR3DC23-Fc_LS for human FcRn was three- to five-fold higher than that of IgG1 isotype and bebtelovimab biosimilar controls with wild type Fc (Fig. 6C and Supplementary fig. 5).

**Figure 6:**
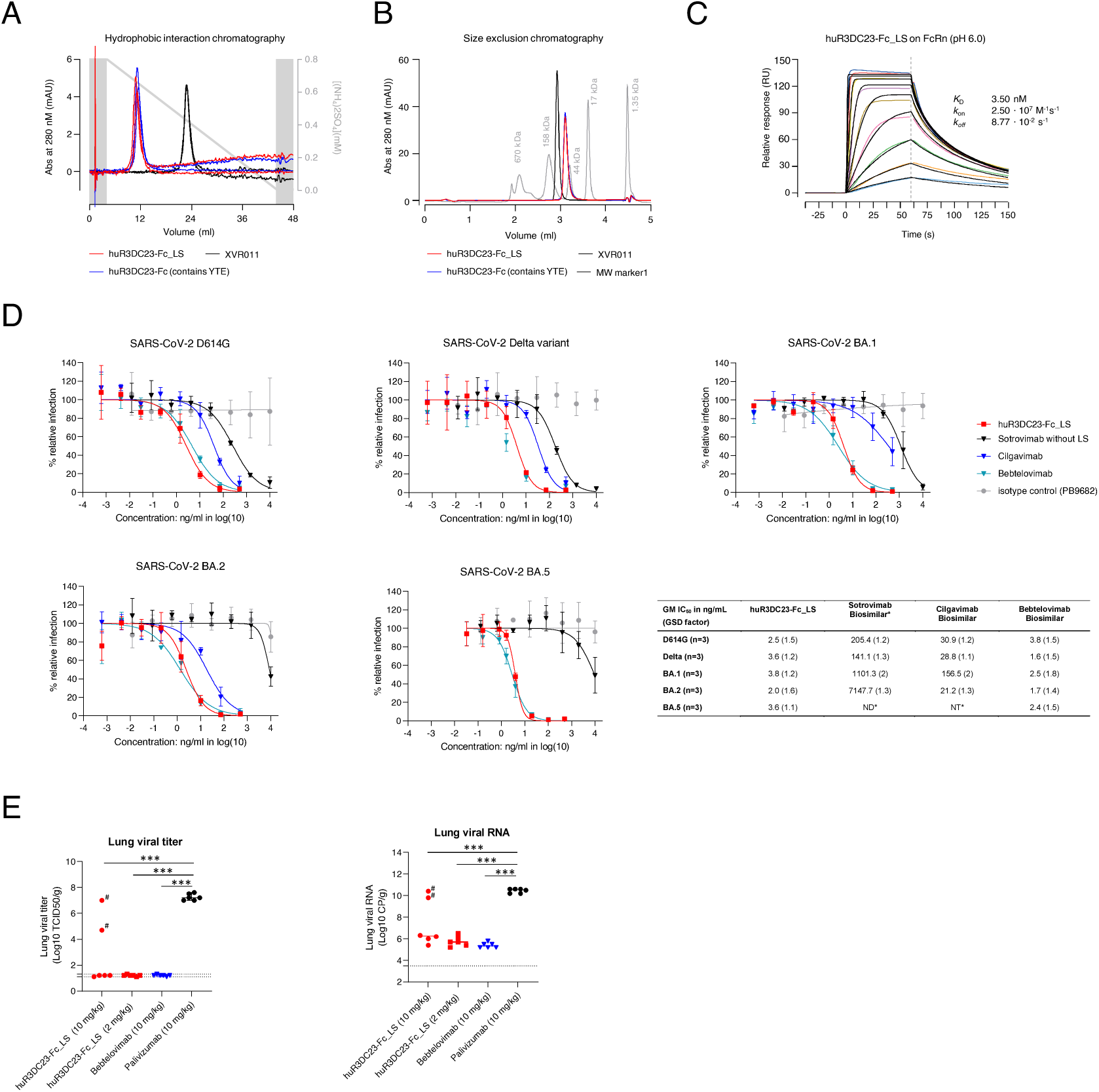
Humanized R3DC23 fused to Fc_LS potently neutralizes a broad range of SARS-CoV-2 variants and controls viral replication in hamsters. **(A)** Analytical hydrophobic interaction chromatography of LS and YTE variants of huR3DC23-Fc overlap. Apparent hydrophobicity was assessed on ProPac HIC-10 HPLC over an (NH_4_)_2_SO_4_ elution gradient (short retention times indicate low apparent hydrophobicity). Duplicate curves are shown for each VHH-Fc. **(B)** Both LS and YTE variants of huR3DC23-Fc elute as a single (overlapping) peak from analytical SEC. Elution profile of molecular weight markers of a gel filtration standard (Bio-Rad) are indicated in grey. Curves and shading indicate mean and SD of triplicate runs. **(C)** FcRn binding of huR3DC23-Fc_LS at pH 6.0 as determined by SPR. After low density immobilization of huR3DC23-Fc_LS to a sensor chip, a 250 to 0.97 nM two-fold dilution series of human FcRn was injected in solution (coloured curves). A 1:1 binding model was fit (black curves). Supporting data in Supplementary figure 5 **(D)** In vitro neutralization of authentic SARS-CoV-2 variants D614G, Delta, Omicron BA.1, Omicron BA.2 and Omicron BA.5 by huR3DC23-Fc_LS as determined in the authentic virus neutralization assay. Data points in the graph represent the mean ± SD (N=3) relative infection. Curves were obtained by non linear regression using the normalized response and only used for visualization purposes. The table shows the geometric mean IC_50_ values (ng/mL) and geometric SD factor (GSD factor) as calculated from 3 independent experiments based on the Zielinska method. (ND*) Not possible to determine IC_50_ value for sotrovimab (without LS) within the tested concentration range (0.128-10 000 ng/ml). (NT*) Cilgavimab was not tested for Omicron BA.5. (*) the IC_50_ of cilgavimab for BA.1 is based on 2 biological replicates in which the highest concentration of cilgavimab (500ng/ml) resulted in a more than 50 % reduction of viral replication but not in a third biological replicate. (Sotrovimab biosimilar*) = sotrovimab biosimilar without LS mutation. **(E)** Treatment with huR3DC23-Fc_LS of SARS-CoV-2 Wuhan infected Syrian Golden hamsters. Male Syrian Golden hamsters were intranasally infected with SARS-CoV-2 (Wuhan strain) on day 0 and received intraperitoneal treatment with either 10 or 2 mg/kg huR3DC23-Fc_LS, 10 mg/kg bebtelovimab (positive control), or 10 mg/kg palivizumab (negative control) 4 hours post-infection. Animals were sacrificed on day 4 and infectious virus (left panel) and viral RNA (right panel) were determined in lung tissue. Horizontal bars indicate the median TCID_50_/gram (left panel) and RNA copies/gram (right panel) lung tissue. Dotted horizontal lines indicate the LLOD. * Two animals in the high dose (10 mg/kg) group were experimentally confirmed to not have been exposed to huR3DC23-Fc_LS treatment. Data were analyzed with the one-way ANOVA and Dunn’s multiple comparison test (***P < 0.0001). (#) Data points corresponding to hamsters for which no or very low levels of huR3DC23-Fc_LS was detected in the serum (Supplementary Figure 6) were omitted from statistical analysis.

huR3DC23-Fc_LS potently neutralized pseudotyped VSV displaying spike proteins of SARS-CoV-2 D614G, BQ.1.1 and XBB with IC_50_ values similar to what was observed for huR3DC23-Fc containing the YTE mutation (Fig. 5D and Table S2). In addition, huR3DC23-Fc_LS potently neutralized VSV pseudotyped with spikes of SARS-CoV-2 BA.4.6, BF.7 and of XBB in which an additional mutation was added, F486P, that results in the higher affinity of the XBB1.5 for ACE2 (XBB.1.5-G252V) (Table S2). In line with what was reported previously, sotrovimab (without LS) and bebtelovimab displayed marked reduction in neutralization activity for XBB and BQ.1.1. (20). Importantly, also authentic SARS-CoV-2 D614G, Delta, BA.1, BA.2 and BA.5 viruses were potently neutralized by huR3DC23-Fc_LS with an IC_50_ of 2.0 to 3.8 ng/ml, close to what was observed for monovalent R3DC23 VHH and huR3DC23-Fc carrying the YTE mutation in virus neutralization assays using authentic D614G and BA.1 SARS-CoV-2 virus (Fig. 6D).

We evaluated the therapeutic potential of huR3DC23-Fc in the Syrian hamster model (21). Hamsters were challenged with an ancestral SARS-CoV-2 isolate (BetaCoV/Munich/BavPat1/2020) and, 4 hours later, treated with either 10 mg/kg or 2 mg/kg huR3DC23-Fc_LS, 10 mg/kg bebtelovimab (biosimilar) or 10 mg/kg palivizumab (negative control treatment) by intraperitoneal injection. At 4 days post infection high levels of huR3DC23-Fc_LS were detected in the serum of all hamsters treated with 2 mg/kg and in 4 out of 6 hamsters treated with 10 mg/kg of this construct. In sharp contrast, no or very low levels of huR3DC23-Fc_LS could be detected in in the sera of two animals that had been treated with 10 mg/kg huR3DC23-Fc_LS (Supplementary fig. 6). This most likely results from unsuccessful injection, which has been observed by others (22). Apart from these 2 hamsters, the lung virus loads, sampled on day 4 after challenge, were below the detection limit in the huR3DC23-Fc_LS treated hamsters whereas control treated animals had significantly higher lung virus loads (Fig. 6 E, left panel). In accordance, in the lungs of hamsters treated with either huR3DC23-Fc_LS or bebtelovimab a strong reduction in viral RNA was observed (Fig. 6E, right panel). This experiment shows that huR3DC23-Fc_LS can strongly restrict SARS-CoV-2 replication in vivo.

## Discussion

It is well established that the level of SARS-CoV-2 neutralizing antibodies is a strong correlate of protection for vaccine efficacy (23). The majority of the neutralizing activity evoked by licensed COVID-19 vaccines is almost exclusively mediated by RBD-specific antibodies (24). Typically, such antibodies can interfere with the interaction of spike trimers with the ACE2 receptor and as such can potently block early events of viral entry. Various RBD-targeting monoclonal antibodies and cocktails thereof have been approved to treat or prevent COVID-19. The importance of these neutralizing antibodies is also highlighted by the continuous emergence of novel SARS-CoV-2 viruses that escape from RBD-targeting antibodies by the acquisition of mutations at sites that are targeted by these antibodies (25). We have isolated a family of single domain antibodies that recognize a quaternary epitope on the conserved membrane proximal region of HR2 and that can neutralize a broad range of SARS-CoV-2 variants with a potency that is surprisingly high for S2-targeting antibodies. We initially tried to determine this epitope via in vitro viral escape mutant selection in the presence of R3DC23, the most potently neutralizing family member, using error prone replication-competent VSV pseudotyped with SARS-CoV-2 spike. The isolated viruses all contained a single substitution at one of 4 different positions (N1192, L1197, L1200 and Q1201) within the membrane-proximal region of HR2. The mutations at position L1197, L1200 and Q1201 resulted in loss of binding and neutralization. However, R3DC23 failed to bind synthetic peptides corresponding to HR2 (1176–1207) comprising the positions that were identified by the viral escape variants (data not shown). Therefore, we probed the R3DC23 binding region on trimeric 2P-stabilized spike ectodomains by HDX-MS. Consistent with the selected escape mutants, binding of R3DC23 protected 2 adjacent peptides from hydrogen-deuterium exchange at the membrane proximal region of HR2. Although HR2 is typically not resolved in structures of full-length spike ectodomain proteins determined by crystallography or cryo-EM, NMR analysis revealed that peptides comprising the full length HR2 of SARS-CoV-1 (which has an identical sequence in SARS-CoV-2) fold as a coiled-coil of 3 parallel HR2 alpha helices (26). To get more detailed insight in the interaction of R3DC23 with the HR2 coiled-coil we determined the crystal structure of R3DC23 in complex with peptides comprising the full length HR2. This revealed that R3DC23 binds to the outer surface of the membrane proximal end of the HR2 coiled-coil at a quaternary epitope comprising two adjacent HR2 helices. This configuration agrees with the mutations that emerged in the viral escape selection. Q1201 is oriented towards the interface of 2 neighboring HR2 helices and is involved in the interaction with the CDR3 via a H-bond network. The leucines at positions 1197 and 1200 that are substituted by prolines in viral escape variants are oriented towards the center of the coiled coil and are not directly involved in the interaction with R3DC23. However, prolines typically act as disrupters of secondary structures and as such typically locate at the ends of alpha helices and are well known to disrupt coiled-coils. As such the L1197P and L1200P substitutions would most likely severely impact the overall conformation of the HR2 coiled coil and hence the quaternary R3DC23 epitope (27). This may explain the attenuated phenotype of these viral escape variants. In contrast, substitution of L1197 and L1200 by respectively phenylalanine and valine or alanine did not affect binding of R3DC23. Although the N1192D escape mutant was selected during replication on VeroE6/TMPRSS2 cells, this mutation did not substantially affect binding of R3DC23 to cells expressing this spike variant nor did it display complete escape from neutralization, especially on VeroE6 cells. This indicates that N1192 is not directly involved in the interaction with R3DC23 but might impact neutralization by this nanobody on VeroE6/TMPRSS2 cells indirectly. Consistently, the escape analysis, HDX-MS and crystal structure indicate that the core of the R3DC23 epitope comprises parts of the N_1194_ESLIDLQE_1202_ HR2 sequence. This sequence is conserved among the SARS-CoV-2 variants of concern or interest that have circulated up to date, explaining the unaffected neutralizing activity of R3DC23 and its Fc-fusions for the SARS-CoV-2 variants tested in this study. During the autumn of 2022 there was a modest and transient circulation of a BA2.75.2 that acquired a D1199N substitution within the HR2 that subsequently reverted to D1199 in the successor CH.1.1 variant. We are currently investigating to which extent this substitution affects R3DC23 binding and neutralization. Although, compared to the rest of the spike stalk, the membrane proximal region of HR2 is relatively conserved between SARS-CoV-2, MERS and HKU-1, the region recognized by R3DC23 contains several differences. One of these is the Q1201K mutation that abrogated the binding of R3DC23 in escape variants. This explains the inability of R3DC23 and the related VHHs to recognize MERS and HKU-1 spikes. In contrast to the HR2 peptide used for crystallization, the HR2 in recombinant S-2P spike that was used for immunization and bio-panning and the spikes on the virion surface that are targeted upon neutralization contain a complex-type glycan at N1194 that is critical for infectivity (28, 29). Although the HR2-targeting nanobodies can efficiently bind to their epitope that is delineated between the viral membrane and this complex N-glycan, this narrow site of approximately 3 nm might be very hard to target by conventional antibodies (17, 30). However, peptide mapping of the spike protein revealed that natural infection with SARS-CoV-2 or even other human coronaviruses can elicits antibodies that can cross-react with a peptide that corresponds to the short flexible linker (K_1205_YEQYIKW_1212_) that sits between the HR2 and the transmembrane region of SARS-CoV-2 spikes (31). Moreover, vaccination of mice with membrane-anchored S2 subunit induced antibodies recognizing this peptide (32). This antibody response might be facilitated by the flexibility of the spike that is provided by flexible linkers within its stem, allowing the spike protein to tilt along its axis to up to 90° (33). This might temporarily expose the flexible linker that separates the HR2 from the transmembrane domain by B cell receptors. It cannot be excluded that the flexibility of the spike protein might also expose the membrane proximal HR2 region below the N1194 glycan. A series of 13 human monoclonal antibodies obtained from a Xenomouse immunized with the SARS-CoV-1 ectodomain have been reported to bind to the HR2 of SARS-CoV-1 (34). The region within HR2 that was recognized by these monoclonal antibodies was, however, not specified. At 25 μg/ml, these antibodies could either partially of completely neutralize lentiviruses pseudotyped with SARS-CoV-1 spikes. Few monoclonal antibodies targeting the N-terminal half of the SARS-CoV-1 HR2, above the N1194 glycosylation site have been described. The humanized antibody (hMab5.17) derived from mice vaccinated with a recombinant bacteria produced non-glycosylated SARS-CoV-1 spike fragment was described to neutralize authentic SARS-CoV-2 with moderate potency (IC_50_ of approximately 12 μg/ml) and to reduce viral replication in Syrian hamsters (35, 36). Similarly, monoclonal antibody 1G10 was obtained from mice vaccinated with a bacteria produced HR2-GST fusion construct. 1G10 could neutralize SARS-CoV-1 with an IC_50_ of 13 µg/ml (37). To the best of our knowledge no potently neutralizing antibody that specifically targets the membrane proximal HR2 region has been reported. It was, however, recently reported that immune serum from mice that had been immunized with a synthetic peptide that corresponds to the carboxy-terminal half of HR2 had modest SARS-CoV-2 neutralizing activity (38). Most likely, the high potency by which R3DC23 can block fusion can be attributed to the quaternary nature of its epitope. Because each R3DC23 VHH binds two adjacent HR2 helices and each HR2 helix in the coiled-coil can be bound by two VHHs, R3DC23 may strongly lock the HR2 coiled-coil. As such, R3DC23 may prevent the unraveling of the HR2 coiled-coil, a critical early step in the spike-controlled membrane fusion process (39, 40). Because R3DC23 neutralizes SARS-CoV-2 via a novel mode of action it was important to test if R3DC23-Fc fusions can also restrict viral replication in vivo. Using the Syrian hamster model we could demonstrate that therapeutic administration of humanized R3DC23 fused to Fc(LS) can effectively restrict viral replication of SARS-CoV-2 in the lungs of infected animals. These results indicate that Fc fusions of R3DC23 either as monotherapy or as part of an antibody cocktail represent a valuable strategy to prevent or treat COVID-19.

## Materials and Methods

### VHH phage display and panning

A llama that was previously immunized with recombinant prefusion stabilized SARS-CoV-1 and MERS spike protein was additionally immunized 3 times, on a weekly basis, with recombinant SARS-CoV-2 spike protein (S-2P) stabilized in its prefusion conformation (13, 14). Five days after the third immunization, peripheral blood lymphocytes were isolated from the llama and a phage display VHH library was constructed. Immunizations and handling of the llama were approved by the Ethical Committee for Animal Experiments of the Vrije Universiteit Brussel (permit No. 16-601-2 granted to VIB Nanobody Core). Phages that display SARS-CoV-2 spike-specific VHHs were enriched from the phage display library by 2 succesive rounds of biopanning on 100 ng of HIS-tagged SARS-CoV-2 spike 2P protein (13), that was immobilized in a well of a microtiter plate (type II, F96 Maxisorp, Nunc) via precoated anti-HIS antibodies in the presence of 10 μg/ml RBD-SD1-mouse IgG. Two additional rounds of biopanning were performed using anti-HIS captured spike proteins (R3C and R4C series) or using directly coated spike proteins (R3DC and R4DC series). These two series of additional rounds of biopanning were also performed in the presence of 10 μg/ml RBD-SD1-mouse IgG. For each panning round an uncoated well was used as a negative control. The wells were then washed 5 times with phosphate-buffered saline (PBS) + 0.05% Tween 20 and blocked with 4% milk powder in PBS (Regilait) in the first panning round and Pierce protein free blocking buffer (Thermo Scientific), SEA blocking buffer (Thermo Scientific) and 1% BSA in PBS in the subsequent panning rounds. Non specifically bound phages were removed by extensive washing with PBS + 0.05% Tween 20. The retained phages were eluted with TEA-solution (14% trimethylamine (Sigma) pH 10) and subsequently neutralized with 1 M Tris-HCl pH 8.0. The collected phages were amplified in exponentially growing E.coli TG1 cells, infected with VCS M13 helper phages and subsequently purified using PEG 8,000/NaCl precipitation for the next round of selection. Enrichment after each panning round was evaluated by infecting TG1 cells with 10-fold serial dilutions of the collected phages after which the bacteria were plated on LB agar plates with 100 μg/mL− ampicillin and 1% glucose.

### Preparation of Periplasmic Extracts

After 3 or 4 panning rounds individual colonies of phage infected bacteria were randomly selected for further analysis. The selected colonies were inoculated in 2 mL of terrific broth (TB) medium with 100 μg/mL ampicillin in 24-well deep well plates. After growing individual colonies for 5 h at 37°C, isopropyl β-D-1-thiogalactopyranoside (IPTG) (1 mM) was added to induce VHH expression during overnight incubation at 37°C. To prepare periplasmic extract, the bacterial cells were pelleted and resuspended in 250 μL TES buffer (0.2 M Tris-HCl pH 8.0, 0.5 mM EDTA, 0.5 M sucrose) and incubated at 41°C for 301min. Subsequently 350 μL water was added to induce an osmotic shock. After 1 h incubation at 4 °C followed by centrifugation, the periplasmic extract was collected.

### Cell Lines

FreeStyle293F cells (ThermoFisher Scientific) and HEK293-S cells (ThermoFisher Scientific) were cultured in FreeStyle293 expression medium (Life Technologies), cultured at 37°C with 8% CO2 while shaking at 130 rpm. HEK293-T cells (ATCC) and Vero E6 cells (ATCC) were cultured at 37°C in the presence of 5% CO2 in DMEM supplemented with 10% heat-inactivated FBS, 1% penicillin, 1% streptomycin, 2 mM l-glutamine, non-essential amino acids (Invitrogen) and 1 mM sodium pyruvate. ExpiCHO-S cells (GIBCO) were cultured at 37°C with 8% CO2 while shaking at 130 rpm in ExpiCHO expression media (GIBCO). Vero E6-TMPRSS2 cells that stably express human TMPRSS2 (NIBIOHN, JCRB1819) (42) were cultured in DMEM containing 10% FBS, Penicillin (100 unit/mL), Streptomycin (100 μg/mL), Geneticin (G418) (1 mg/ml). Vero E6-TMPRSS2 cells were seeded in medium without Geneticin for infectivity assays.

### Production of VHHs in E. coli

For the production of VHH in E. coli, a pMECS vector containing the VHH of interest was transformed into WK6 cells (the non-suppressor E coli strain) and plated on an LB plate containing ampicillin. The next day clones were picked and grown overnight in 2mL LB containing 100 μg/ml ampicillin and 1% glucose at 37°C while shaking at 200 rpm. One ml of this preculture was used to inoculate 25 ml of TB supplemented with 100 µg/ml ampicillin, 2mM MgCl2 and 0.1% glucose and incubated at 37°C with shaking (200-250 rpm) untill an OD600 of 0.6-0.9 was reached. VHH expression was induced by addition of IPTG to a final concentration of 1mM followed by overnight incubation at 28°C while shaking at 200 rpm. The VHH-containing fraction was extracted from the periplasm and purified as described in Wrapp et al. (14). In short, the VHHs were purified from the periplasmic extract using Ni Sepharose beads (GE Healthcare). After elution with 500 mM imidazole the VHH-containing fractions were buffer-exchanged with PBS using a Vivaspin column (5 kDa cutoff, GE Healthcare). The purified VHHs were analyzed by SDS-PAGE and Coomassie staining and by intact mass spectrometry. For crystallography, R3DC23 was produced in a 1L culture of E. coli WK6. The periplasmic extract was loaded on a 1 ml HisTrap^TM^ High Performance column (Cytiva), which had been equilibrated with 20 mM NaH_2_PO_4_ pH 7.5, 0.5 M NaCl, and 20 mM imidazole. Bound VHH was eluted with 20 mM NaH2PO4, 20 mM NaCl and 0.5 M imidazole. The peak fractions were pooled and further purified by size exclusion chromatography, using a HiLoad 16/600 Superdex® 75 prep grade column (Sigma-Aldrich) and were eluted in PBS.

### Production of YTE variants of VHH-Fc in mammalian cells

HEK293S cells were transfected with VHH-Fc (S) encoding plasmids using poly-ethylene immine (PE)I. Briefly, suspension-adapted and serum-free HEK293S cells were seeded at 3 × 106 cells/mL in Freestyle-293 medium (ThermoFisher Scientific). Next, 4.5 μg of pcDNA3.3-VHH-Fc plasmid DNA was added to the cells and the cells were incubated on a shaking platform at 37°C and 8% CO2, for 5 min. Next, 9 μg of PEI was added to the cultures, and cells were further incubated for 5 h, after which an equal culture volume of Ex-Cell-293 (Sigma) was added to the cells. Transfected cells were incubated for 4 days, after which the cells were pelleted (10 min, 300 g) and supernatants were filtered before further use.

Additionally, huR3DC23-Fc_YTE was expressed in ExpiCHO-STM cells (ThermoFisher Scientific), according to the manufacturer’s protocol. Briefly, a 50 mL culture of 6 x 106 cells per mL, grown at 37°C and 8% CO2, was transfected with 40 µg of pcDNA3.3-VHH72-Fc plasmid DNA using of ExpiFectamineTM CHO reagent. One day after transfection, 300 µL of ExpiCHOTM enhancer and 8 mL ExpiCHOTM feed was added to the cells, and cultures were further incubated at 32°C and 5% CO2. Cells were fed a second time on day 5 after transfection. Productions were collected as soon as cell viability dropped below 75%.

For purification of the VHH-Fc proteins, supernatants were loaded on a 5 mL MAbSelect SuRe column (Cytiva). Unbound proteins were removed by a wash step with McIlvaine buffer pH 7.2, and bound proteins were eluted using McIlvaine buffer pH 3. Immediately after elution, protein-containing fractions were neutralized with 30% (v/v) of a saturated Na_3_PO_4_ buffer. Next, these fractions were pooled, and loaded on a HiPrep Desalting column for buffer exchange to PBS pH7.4.

### Acquisition of sotrovimab (without LS), cilgavimab, bebtelovimab biosimilars and palivizumab

Bebtelovimab biosimilar (PX-TA1750), cilgavimab biosimilar (PX-TA1033) and sotrovimab biosimilar (without LS) (PX-TA1637) were commercially purchased from Proteogenix. Clinical grade palivizumab was obtained from the Ghent University hospital.

### Production of huR3DC23-Fc_LS following transient transfection

The gene encoding huR3DC23-Fc_LS was codon optimized, synthesized, and cloned into the pXLG6 backbone vector at ATUM’s laboratories. Upon gene and codon optimization the R3DC23 DNA sequence was inserted into pXLG6 expression vector and transfected in CHOExpress ™ cells at a cell density of 4.00E+6 cells/ml. TGE supernatant was harvested by centrifugation and clarified by filtration (0.2µm) after 10 days when cell viability dropped below 90%. The protein was further purified by Protein A.

### Fed batch production of huR3DC23-Fc_LS from stable pool at 1L scale

The gene encoding huR3DC23-Fc_LS was codon optimized, synthesized, and cloned into the pXLG6 backbone vector at ATUM’s laboratories. Upon expansion to a density of about 4 x 10^6^ cells/ml, parental CHOExpress™ cells were co-transfected with the expression vector and the pXLG5 helper vector. The stable pool was generated under 50 mg/L puromycin selective pressure (applied daily) and further expanded. The stable pool research cell bank was banked at day 14 when cell viability reached 95%.

The RCB pool was then expanded for protein production at 1L scale and cultured until day 12 (cell density 3.5 x 10^7^ cells/mL, cell viability 96%). The supernatant was harvested by centrifugation and clarified by filtration (0.2 µm). The protein was further purified by Protein A using MabSelect SuRe LX resin. Consecutive washed were performed with 20 mM sodium phosphate and 110 mM NaCl at pH 7.2; 100 mM sodium acetate and 500 mM NaCl at pH 5.5; and 20 mM sodium phosphate at pH 7.2. The eluate in 100 mM sodium acetate pH 3.5 was neutralized to pH 7.0 by addition of 1 M Tris pH 11.0 (10%_v/v_). After filter sterilization (0.22 µm), the protein was aliquoted at 2 mg/ml.

### Protein preparation for biophysical analysis

Preceding biophysical characterization, MAbSelect SuRe-purified protein samples were further purified via size-exclusion chromatography (SEC) on a 12 °C-cooled Superdex 200 column (Cytiva) equilibrated with either phosphate-buffered saline (PBS, Sigma-Aldrich) supplemented with 0.02% sodium azide to prevent microbial growth, or a sample buffer comprising 50 mM L-histidine and 150 mM L-arginine (Sigma-Aldrich), 0.02% polysorbate-20 and 0.02% sodium azide, set to pH 7.0 at 25 °C. After filter sterilization (0.22 µm), 1 mg/ml aliquots were snap-frozen in polypropylene tubes in liquid nitrogen and stored at -80 °C.

### Sample composition

Purified VHH-Fc samples were characterized by analytical SEC to determine the molecular composition of each sample. After rapid thawing in a water bath at 25 °C, 10 min centrifugation at 16,000 xg and transfer of supernatant to fresh tubes, 5 µg was injected on an AdvanceBio SEC column, 4.6 x 300 mm (Agilent) with 2.7 µm porous particle size and 300 Å pore size, calibrated with PBS. The separation was monitored by absorbance at 280 nm with a 16 nm bandwidth, without reference subtraction. For additional quality control, proteins were separated on reducing 15% SDS-PAGE with Coomassie staining.

### Hydrophobic interaction chromatography (HIC) assay

Apparent hydrophobicity was assessed using a hydrophobic interaction chromatography (HIC) assay employing a Dionex ProPac HIC-10 column, 100 mm × 4.6 mm (Thermo Fisher 063655), containing a stationary phase consisting of a mixed population of ethyl and amide functional groups bonded to silica. All separations were carried out on an Agilent 1100/1260 HPLC equipped with a UV/VIS detector. The column temperature was maintained at 25 °C throughout the run and the flow rate was 0.8 ml/min. The mobile phases used for HIC were 1.6 M ammonium sulfate and 50 mM phosphate pH 7.0 (buffer A-, and 50 mM phosphate pH 7.0 (buffer B). Protein and calibrator samples were diluted 1:1 with buffer A and injected onto the column. Following a 5 min hold at 50% B, bound protein was eluted using a linear gradient from 50 to 100% B in 50 min followed by 5 min hold at 100% B. The column was washed with 100% B, followed by 50 mM ammonium acetate pH 5.0 and re-equilibration in 50% B for 10 min prior to the next sample. The separation was monitored by absorbance at 280 nm with a 16 nm bandwidth, without reference subtraction.

### Mass spectrometry analysis of proteins

Intact VHH was separated on an Ultimate 3000 HPLC system (Thermo Fisher Scientific, Bremen, Germany) online connected to an LTQ Orbitrap Elite mass spectrometer (Thermo Fischer Scientific). Briefly, approximately 2 µg of protein was injected on a Zorbax Poroshell 300SB-C8 column (5 µm, 300Å, 1×75mm IDxL; Agilent Technologies) and separated using a 15 min gradient from 5% to 80% solvent B at a flow rate of 100 µl/min (solvent A: 0.1% formic acid and 0.05% trifluoroacetic acid in water; solvent B: 0.1% formic acid and 0.05% trifluoroacetic acid in acetonitrile). The column temperature was maintained at 60°C. Eluting proteins were directly sprayed in the mass spectrometer with an ESI source using the following parameters: spray voltage of 4.2 kV, surface induced dissociation of 30 V, capillary temperature of 325 °C, capillary voltage of 35 V and a sheath gas flow rate of 7 (arbitrary units). The mass spectrometer was operated in MS1 mode using the orbitrap analyzer at a resolution of 120,000 (at m/z 400) and a mass range of 600-4000 m/z, in profile mode. The resulting MS spectra were deconvoluted with the BioPharma FinderTM 3.0 software (Thermo Fischer Scientific) using the Xtract deconvolution algorithm (isotopically resolved spectra). The deconvoluted spectra were manually annotated.

### Enzyme-linked immunosorbent assay

Wells of microtiter plates (type II, F96 Maxisorp, Nunc) were coated overnight at 4°C with 100 ng of recombinant SARS-Cov-2 S-2P, SARS-CoV-2 S-6P protein, recombinant SARS-CoV-1 spike protein, recombinant MERS-CoV spike protein, recombinant HKU1 spike protein (all kind gifts by Dr. Jason Mclellan) SARS-CoV-2 Omicron BA.1 S protein (ACROBiosystems), mouse Fc-tagged SARS-CoV-2 RBD (Sino Biological, 40592-V05H), the SARS-CoV-2 spike S2 subunit (ACROBiosystems, S2NC52H5) or BSA. The coated plates were blocked with 5% milk powder in PBS. Dilution series of the VHHs, VHH-Fcs or antibodies were added to the wells and plates were further incubated for 90 minutes at room temperature. After washing, binding of VHHs, was detected by HRP-conjugated rabbit anti-camelid VHH antibodies (Genscript, A01861-200, 1/5000) or a mouse anti-HA antibody (BioLegend 901501, 1/2000), followed by anti-mouse IgG-HRP (Cytiva, NA931V, 1/2000). Binding of VHH-Fcs or conventional human monoclonal antibodies was detected by HRP-conjugated rabbit anti-human IgG (Sigma, A8792, 1/2000). After washing 50 μL of TMB substrate (Tetramethylbenzidine, BD OptEIA) was added to the plates and the reaction was stopped by addition of 50 μL of 1 M H_2_SO4. The absorbance at 450 nm was measured with an iMark Microplate Absorbance Reader (Bio Rad). Curve fitting was performed using nonlinear regression (Graphpad 8.0).

### Flow cytometric analysis of binding to HEK293T cells expressing the SARS-CoV spike protein

Binding of VHHs to spike proteins on the surface of mammalian cells was determined by flow cytometry using pcG1-expression plasmids containing the coding sequence of the SARS-CoV-2 spike protein from which the C-terminal 18 amino acids were deleted and in which the D614G substitution was introduced by QuickChange site-directed mutagenesis (Agilent) according to the manufacturer’s instructions. Two days after transfecting HEK293T cells with a GFP expression plasmid in combination with either a spike expression plasmid or a control expression plasmid the cells were collected. All further procedures were performed on ice. The cells were washed once with PBS and blocked with 1% BSA. The cells were then stained with antibody or VHH dilution series for 90 minutes and subsequently washed 3 times with PBS containing 1% BSA. Binding of VHHs was detected using a mouse anti-HIS-tag antibody (Biorad, MCA1396, 1/2000) and an AF647 conjugated donkey anti mouse IgG antibody (Invitrogen, A31571, 1/1000). Binding of VHH-Fcs or antibodies was detected using an AF633 donkey goat anti-human IgG antibody (Iinvitrogen, A21091, 1/1000) and dead cells were stained using Live/Dead stain (Invitrogen, 15560607, 1/1000). Following 3 washes with PBS containing 0.5% BSA, the cells were analyzed by flow cytometry using a BD LSRII flow cytometer (BD Biosciences). The binding curves were fitted using nonlinear regression (Graphpad 9.10).

### Pseudovirus neutralization assay with periplasmic extracts

Pseudoviruses expressing the SARS-CoV-2 spike (D614G) were incubated for 30 min at 37°C with a 1/100 dilution of periplasmic extract in Fluorobrite DMEM medium (Invitrogen), supplemented with 5% heat-inactivated FBS, 1% penicillin, 1% streptomycin, 2 mM L-glutamine, non-essential amino acids (Invitrogen) and 1 mM sodium pyruvate. The incubated pseudoviruses were subsequently added to subconfluent monolayers of Vero E6 cells from which the original growth medium was removed. Sixteen hours later, the cells were lysed using passive lysis buffer (Promega). The transduction efficiency was quantified by measuring the GFP fluorescence in the prepared cell lysates using a Tecan infinite 200 pro plate reader. GFP fluorescence was normalized using the GFP fluorescence of non-infected cells and infected cells treated with PBS.

### FcRn binding affinity

Surface plasmon resonance (SPR) analysis of huR3DC23-Fc_LS binding to purified recombinant human FcRn / FCGRT-B2M protein was performed by FairJourney Biologics (Porto, Portugal) on a Biacore 8 K+ instrument. His-tagged recombinant human FcRn (FCGRT-B2M heterodimer) protein was purchased from ACROBiosystems. Bebtelovimab biosimilar and a human IgG1 isotype control antibody were used as controls in the assay. In brief, the experimental set-up was as follows: huR3DC23-Fc_LS or control antibodies were immobilized at low density on a CM5 sensor chip (Cytiva) by amine coupling to immobilization levels of 130 to 283 RU. At pH 6.0, human FcRn was injected in solution at 1.5 µM (anchor point) and in 2-fold dilution series in the range of either 1000 to 7.8 nM (control antibody-immobilized channels), or 250 to 0.97 nM (huR3DC23-Fc_LS-immobilized channels). At pH 7.4, human FcRn was injected in solution at 1.5 µM (anchor point) and in 2-fold dilution series in the range of 1000 to 7.8 nM for all immobilized channels. Analyte injections were performed in running buffer during 60 seconds at 30 µl/min; assay runs with 0 nM concentration were included as blank reference. SPR running buffer consisted of PBS with 0.05% Tween 20 at pH 6.0 or 7.4. A multicycle kinetics protocol was applied (off-rate measurement 90 seconds); data obtained from antibody or VHH-Fc concentrations above 250 nM was omitted because of bulk effects. After double reference subtraction, data was analyzed either using the steady state affinity predefined evaluation method of the Biacore Insight Evaluation Software, or by fitting a 1:1 binding model in the same software.

### SARS-CoV pseudovirus neutralization assay

To generate replication-deficient VSV pseudotyped viruses, HEK293T cells, transfected with SARS-CoV-1 S or SARS-CoV-2 S expression vectors were inoculated with a replication-deficient VSV vector containing eGFP and firefly luciferase expression cassettes (43, 44). After 1 h incubation at 37°C, the inoculum was removed, cells were washed with PBS and incubated in media supplemented with an anti-VSV G mAb (ATCC CRL-2700, RRID:CVCL_G654) for 16 hours. Pseudotyped particles were then harvested and clarified by centrifugation (44). For the VSV pseudotype neutralization experiments, the pseudoviruses were incubated for 30 min at 37°C with different dilutions of purified VHH or VHH-Fc fusions or with GFP-binding protein (GBP: a VHH specific for GFP). The incubated pseudoviruses were subsequently added to subconfluent monolayers of Vero E6 or VeroE6/TMPRSS2 cells. Sixteen hours later, the cells were lysed using passive lysis buffer (Promega). The transduction efficiency was quantified by measuring the GFP fluorescence in the prepared cell lysates using a Tecan infinite 200 pro plate reader. GFP fluorescence was normalized using either the GFP fluorescence of non-infected cells and infected cells treated with PBS or the lowest and highest GFP fluorescence value of each dilution series. The IC_50_ was calculated by non-linear regression curve fitting, log(inhibitor) vs. response (four parameters). Alternatively, dilution series of VHHs or antibodies were mixed with 100 PFU of GFP-expressing replication competent VSV virus particles pseudotyped with the SARS-CoV-2 spike protein derived from an early isolate. Of note during propagation of this viral clone (S1-10a) on Vero E6 cells the furin cleavage site was mutated and as such inactivated (45). After 30 minutes incubation at 37°C the virus-antibody mixtures were added to monolayers of Vero E6 cells and allowed to infect and replicate for three days. In all neutralization assays using pseudotyped VSV viral particles FluoroBrite DMEM medium (Invitrogen) supplemented with 5% heat-inactivated FBS, 100 unit/mL penicillin, 100 unit/mL streptomycin, 2 mM L-glutamine, non-essential amino acids (Invitrogen) and 1 mM sodium pyruvate was used to prepare the VHH or antibody-virus mixtures. The mixtures were added to cells from which the original growth medium was removed. For all VSV pseudotype neutralization assays using huR3DC23-Fc_LS, huR3DC23-Fc_LS from stable cell pools was used.

### Neutralization assays using authentic SARS-CoV-2 performed at VIB-UGent CMB

In vitro neutralization experiments with authentic SARS-CoV-2 viruses were performed in the Biosafety level 3 laboratory at VIB-UGent Center for Medical Biotechnology. The plaque reduction assays using authentic viruses were performed with SARS-CoV-2 D614G strain SARS-CoV-2/human/FRA/702/2020, obtained from the European Virus Archive (EVAG) and with an SARS-CoV-2 BA.1 virus that was obtained from KULeuven (46) and grown at KULeuven on Vero E6 cells. Further propagation of the virus was performed on VeroE6/TMPRSS2 cells. Both viruses were titrated using a plaque assay in which monolayers of VeroE6/TMPRSS2 cells were infected with dilutions series prepared in Dulbecco’s Modified Eagle Medium (DMEM) supplemented with 2% fetal bovine serum (FBS) in duplicate for 2 vials of each virus. Two hours after infection, Avicel was added to a final concentration of 0.3% (w/v). Dose-dependent neutralization was assessed by mixing the constructs at different concentrations (5-fold serial dilutions) with 40 PFU of SARS-CoV-2, followed by incubation of the mixture at 37°C for 1 hour. The VHH-virus mixes were then added to VeroE6/TMPRSS2 cell monolayers in 24-well plates and incubated at 37°C for 1 hour. Subsequently, Avicel was added to a final concentration of 0.3% (w/v). After 2 days of incubation at 37°C, the overlays were removed, and the cells were fixed with 3.7% paraformaldehyde (PFA) and stained with 0.5% crystal violet. Half-maximum neutralization titers (PRNT_50_) were defined as the VHH or VHH-Fc concentration that resulted in a plaque reduction of 50% across two independent plates.

### Neutralization assays using authentic SARS-CoV-2 to test the neutralizing activity of huR3DC23-Fc_LS

SARS-CoV-2 viruses belonging to different lineages (D614G, Delta, Omicron BA.1, Omicron BA.2 and Omicron BA.5) were isolated from nasopharyngeal swabs taken from patients/travelers between January 2020 and July 2022. More specifically, the following clinical isolates were used: SARS-CoV-2 Isolate BavPat1/2020/Germany (09 Feb 2020); Delta variant SARS-Related Coronavirus 2, Isolate hCoV-19/USA/MD-HP05647/2021; Omicron BA.1 variant SARS-CoV-2 hCoV-19/Netherlands/NH-RIVM-72291/2021; Omicron BA.2 variant Clinical isolate hCoV-19/Netherlands/VCB-20220303-1/2022; and Omicron BA.5 variant Clinical isolate hCov19/NL/VCB-20220714-2/2022. Dose dependent neutralization of the test item (huR3DC23-Fc_LS), the positive controls (bebtelovimab biosimilar, cilgavimab biosimilar, sotrovimab biosimilar (without LS) and a negative control (isotype control) were assessed in an authentic virus neutralization assay. For all assays in which authentic D614G, Delta, BA.1, and BA.2 SARS-CoV-2 were tested, huR3DC23-Fc_LS produced from transiently transfected cells was used. For all assays in which SARS-CoV-2 BA.5 was tested, huR3DC23_Fc_LS produced from stable cell pools was used. For each variant, three independent runs were performed. Different system controls were included in the assay: cell only (medium only), virus only, and an internal positive control (human serum). Briefly, 5-fold or 7-fold serial dilutions of the test items and controls were incubated with a fixed amount of plaque-forming units (PFUs) of the virus for 1 hour at room temperature. Afterwards, the Vero E6 cell monolayer was inoculated with virus antibody mixtures for 1 hour at 37°C. In a next step, the inoculum was removed and cells were incubated at 37°C with infection medium (up to 18-24 hours post-infection). Afterwards, the SARS-CoV2 infected cells were fixed and immunostained with a SARS-CoV Nucleocapsid Antibody (Sino Biological, Catalogue number: 40143-MM05), followed by HRP-conjugated Goat anti-Mouse IgG (H+L) Secondary Antibody (Invitrogen, catalogue number A16072). Spots (infected cells) were counted using an Immunospot Image Analyser. For each test item, the compound concentration showing 50 % reduction in infection (IC_50_) was calculated based on the Zielinska method. The geometric mean values were calculated based on three independent runs.

### S1 shedding assay

Antibody or VHH was added at a final concentration of 10 µg/ml to 500 000 HEK293T cells transfected with a SARS-CoV-2 spike (D614G)-expressing plasmid, or an empty control vector. The antibody-cell mixture was incubated for 30 min at 37°C and 5% CO_2_. After incubation, cells were pelleted by centrifugation, supernatant was transferred to a fresh tube and the cell pellet was lysed with 250 µl of RIPA lysis buffer (50 mM Tris-HCl pH 8.0, 100 mM NaCl, 1mM EDTA, 1mM EGTA, 0.1% SDS, 1% NP-40). Twenty µl samples of supernatant and lysate were separated on 8% SDS-PAGE gels, and electroblotted onto nitrocellulose membranes. Membranes were blocked with 4% milk, stained with rabbit anti-SARS-S1 antibody (1/1000, Sino Biological, 40591-T62) followed by anti-rabbit IgG-HRP (1/2000, GE Healthcare, NA934V) and developed using Pierce™ ECL Western Blotting Substrate (Thermofisher Scientific).

### Fusion inhibition assay using replication-competent GFP report VSV virus pseudotyped with Wuhan SARS-CoV spikes(del-18)

VeroE6/TMPRSS2 cells were infected with 40 PFU of SARS-CoV-2 spike GFP-expressing pseudotyped replication-competent VSV-GFP virus, a generously provided by Dr. Florian Schmidt (45). Two hours, later the indicated monoclonal antibodies or VHHs were added. Non-infected cells were used as negative controls. Infected cells were incubated overnight and imaged with a fluorescence microscope. GFP fluorescence was measured with a fluorimeter. Of note, different from the clone of replication-competent pseudotyped VSV particles used in the neutralization assays, the furin cleavage site of the virus used in the fusion assays had an intact furin cleavage site as confirmed by Sanger sequencing.

### Fusion inhibition assay using spike-expressing Vero E6 cells

Vero E6 cells were transfected with an GFP expression vector together with either a control expression vector (no spike) or a SARS-CoV-2 spike expression vector using Fugene (Promega). Two hours after transfection, PBS, monoclonal antibodies, or VHHs were added to a final concentration of 10 μg/ml. Twenty-two or forty hours later, the cells were imaged using an Olympus fluorescence microscope using a 10x lens. Alternatively, monoclonal antibodies or VHHs were added to transfected cells at a final concentration of 1 µg/ml and GFP expression was monitored hourly over time with an Incucyte Zoom live cell analysis device (Sartorius) and analyzed with the provided Incucyte software.

### Viral escape selection

Monolayers of VeroE6/TMPRSS2 cells seeded in 96 well plates were infected with 200 PFU of GFP expressing replication-competent VSV virus particles pseudotyped with the SARS-CoV-2 spike protein containing an intact furin cleavage site, generously provided by Dr. Florian Schmidt (45). Two hours after infection, 10 μg/ml of VHH R3DC23 was added and control wells without VHH were included. From the growth medium of wells that displayed syncytia formation or viral replication in the presence of VHH R3DC23, single clones were isolated by limiting dilution on fresh VeroE6/TMPRSS2 cells seeded in 96 well plates. Growth medium of wells with a single PFU was used to propagate possible escape viruses on monolayers of VeroE6/TMPRSS2 cells seeded in 6 well plates in the presence of VHH R3DC23. From these infected cells, RNA was prepared using a nucleospin RNA virus kit (Macherey Nagel Bioanalysis) and used to generate cDNA using random hexamer primers. This cDNA was used to amplify the spike coding sequences by PCR. The resulting PCR fragments were purified and sequenced using Sanger sequencing. The obtained nucleotide sequences were analyzed and aligned to spike proteins of WT SARS-CoV-2 and clade 1, 2, and 3 sarbecoviruses (47) using CLC Main Workbench 20.0.4. Mutations were visualized on a model of full-length glycosylated spike protein obtained from Charmm-gui.org (PDB: 6VXX_1_1_1 model) or the SARS-CoV-2 HR2 coiled-coil as determined by NMR (PDB: 3FXP) using Pymol.

### Growth kinetics of viral escape variants

Vero E6 cells seeded in a 96 well-plate were infected with 50 PFU of GFP-expressing replication competent VSV virus particles pseudotyped with the SARS-CoV-2 spike protein, that were obtained during escape selection. GFP-expression was monitored hourly with an Incucyte Zoom device and analyzed with accompanying software.

### HR2 expression and purification

For structural biology purposes the HR2 protein was expressed in a bacterial expression system. Therefore, the synthetic gene encoding residues H1159-K1211 of the SARS-CoV-2 spike was cloned into a pFloat-SUMO vector, generating a His-tagged SUMO-HR2 fusion protein. The construct also contained a 3C protease cleavage site to remove the His-SUMO-tag. The pFloat-SUMO-HR2 plasmid was transformed in BL21(DE3) cells and plated on kanamycin (100 µg/ml) containing LB agar plates. A small LB culture, supplemented with 100 µg/ml kanamycin, was inoculated with a single colony of BL21(DE3)(pFloat-SUMO-HR2) and grown overnight at 37°C. 1L LB cultures were subsequently inoculated with 20 ml of this preculture and grown at 37°C until OD_600_ reached 0.8. At this point protein expression was induced by adding 0.5 mM isopropyl β-D-1-thiogalactopyranoside (IPTG) to the E. coli culture. Cells were incubated further overnight at 20°C and subsequently harvested by centrifugation (Beckman rotor 8.1000, 5000 rpm, 15 min, 4°C). The pellet was resuspended in PBS, 500 mM NaCl, 10 mM imidazole, 5 mM ß-mercaptoethanol, 0.1 mg/mL 4-(2-aminoethyl) benzenesulfonyl fluoride hydrochloride (AEBSF), 1 µg/mL leupeptine, 50 µg/mL DNaseI and 20 mM MgCl_2_. The cells were lysed using a French press (Constant Systems) at 20 kpsi and the cell debris was removed by centrifugation. The cell lysate was loaded on a Ni-sepharose FF HiLoad column (GE Healthcare), equilibrated in 20 mM Tris-HCl pH 7.5, 500 mM NaCl, 10 mM imidazole, 5 mM ß-mercaptoethanol. The bound proteins were eluted using a linear gradient to 500 mM imidazole. Fractions containing the His-SUMO-HR2 protein were pooled and dialysed overnight to 20 mM Tris-HCl pH 7.5, 150 mM NaCl at 4°C, followed by 2h incubation with 3C protease at room temperature. The cleaved sample was loaded again on a Ni-sepharose FF HiLoad column, equilibrated in the same buffer. The flow through, containing the HR2 protein, was concentrated and applied to a BioRad Enrich70 10/30 size exclusion column (SEC), equilibrated in 20 mM Tris-HCl pH 7.5, 150 mM NaCl. The HR2-containing SEC fractions were pooled.

### R3DC23-HR2 crystallization, X-ray data collection, processing, and structure determination

For crystallization, R3DC23 was added to HR2 in 1.2 times molar excess and concentrated to 19 mg/ml using Amicon Ultra 3 kDa cut off centrifugal filter devices. Crystallization screens were set up using the sitting drop vapor diffusion technique, mixing 0.1 µl of R3DC23-HR2 and 0.2 µl bottom solution. Crystals were grown from the Molecular Dimensions Proplex crystallization screen, in 0.1 M magnesium chloride hexahydrate, 0.1 M sodium citrate pH 5.0, and 15% PEG4000. For X-ray data collection, crystals were flash frozen in liquid nitrogen. X-ray data were collected on the i24 beamline at the Diamond Light Source synchrotron facility (Didcot, UK). X-ray data were processed using autoPROC+Staraniso (48, 49). The structure of the R3DC23-HR2 complex was solved using the automatic molecular replacement workflow in the CCP4 cloud (50). The initial model was further build manually in Coot (51) and refined using phenix.refine (52) from the Phenix crystallographic software suite (53). Data collection parameters, as well as processing and refinement statistics are shown in Table S1.

### Generation of R3DC23-Fc(YTE)

A humanized (Q1D, Q5V, A13P, D15G, T19R, M63V, S73N, T84L, K83R and Q98L; according to Kabat numbering) version of R3DC23 was fused via a (G4S)_2_ linker to a human IgG1 Fc (EPKSCdel_YTE_K447del) and ordered synthetically at IDT as gBlocks. Upon arrival, gBlocks were solubilized in ultraclean water at a concentration of 20 ng/µL. gBlocks were A-tailed using the NEBNext-dA-tailing module (NEB), purified using CleanPCR magnetic beads (CleanNA) and inserted in pcDNA3.4-TOPO vector (ThermoFisher). The ORF of positive clones was fully sequenced, and pDNA of selected clones was prepared using the NucleoBond Xtra Midi kit (Machery-Nagel).

### Generation of spike protein expression vectors for the production of VSVdelG pseudovirus particles expressing spike proteins of SARS-CoV-2 variants

The pCG1 expression vector for the SARS-CoV-2 spike protein containing the D614G mutation was generated from the pCG1-SARS-2-Sdel18 vector by introducing the specific RBD mutation(s) via QuickChange mutagenesis using appropriate primers, according to the manufacturer’s instructions (Aligent). For the pCG1-SARS-2-Sdel18 expression vector for the omicron BA.1 variant, a codon optimized spike protein nucleotide sequence containing the BA.1 mutations (A67V, Δ69-70, T95I, G142D, Δ143-145, N211I, Δ212, ins215EPE, G339D, S371L, S373P, S375F, K417N, N440K, G446S, S477N, T478K, E484A, Q493R, G496S, Q498R, N501Y, Y505H, T547K, D614G, H655Y, N679K, P681H, N764K, D796Y, N856K, Q954H, N969K, L981F) and flanking BamHI and SalI restriction sites was ordered at GeneArt (Thermo Fischer Scientific) and cloned in the pCG1 vector as an BamHI/SalI fragment. For the pCG1-SARS-2-BA.2 Sdel18 expression vector, a codon-optimized spike protein nucleotide sequence containing the BA.2 mutations (T19I, ΔL14-P26, A27S, G142D, V213G, G339D, S371F, S373P, S375F, T376A, D405N, R408S, K417N, N440K, S477N, T478K, E484A, Q493R, Q498R, N501Y, Y505H, D614G, H655Y, N679K, P681H, N764K, D796Y, Q954H, N969K) and flanking BamHI and SalI restriction sites was ordered at GeneArt (Thermo Fischer Scientific) and cloned in the pCG1 vector as an BamHI/SalI fragment. After sequencing, clones containing the correct spike coding sequence were prepared using the Qiagen plasmid Qiagen kit. For the pCG1-SARS-2-BA.2.75 Sdel18 expression vector, a codon-optimized spike protein nucleotide sequence containing the BA.2 mutations (T19I, ΔL14-P26, A27S, G142D, S147E, W152R, F157L, I210V, V213G, G257S, G339D, S371F, S373P, S375F, T376A, D405N, R408S, K417N, N440K, G446S, S477N, N460K, T478K, E484A, Q498R, N501Y, Y505H, D614G, H655Y, N679K, P681H, N764K, D796Y, Q954H, N969K) and flanking BamHI and SalI restriction sites was ordered at GeneArt (Thermo Fischer Scientific) and cloned in the pCG1 vector as an BamHI/SalI fragment. To generate expression plasmids for the BA.5 spike protein mutations ΔLH96-V70, L452R, F486V and R493Q were introduced in the BA.2 spike expression construct by QuickChange mutagenesis according to the manufacturer’s instructions (Aligent). Likewise, to generate the BA.4.6 and BQ.1.1 spike expression vectors mutations R346T and N658S and mutations R346T and K444T were respectively introduced in the BA.5 spike expression vector by QuickChange mutagenesis. The BF.7 spike expression vector was generated by introducing the R346T substitution in the BA.5 spike expression vector by QuickChange mutagenesis. After sequencing, plasmids from clones containing the correct spike coding sequence were prepared using the Qiagen plasmid Qiagen kit. To generate the pCG1-SARS-2-XBB, a Gblock corresponding to the SARS-CoV-2 amino acid sequence V70-V510 containing the XBB mutations (V83A, G142D, ΔY144, H146Q, Q183E, V213E, D339H, R346T, L368I, S371F, S373P, S375F, T376A, D405N, R408S, K417N, N440K, V445P, G446S, S477N, T478K, E484A, F486S, F490S, Q498R, N501Y, Y505H) was ordered and cloned into the pCG1-SARS-2BA.2 vector via Gibson assembly according to the manufacturer’s instructions (New England Biolabs). To this end the pCG1-SARS-2-BA.2 vector was amplified by PCR using appropriate primers (GTGCCATTGGTGCCGGACACG and CACCAGCCTTACAGAGTGG) and the Phusion High-Fidelity DNA polymerase (New England Biolabs). The XBB.1.5(-G252V) spike expression vector was generated by introducing the F496P substitutions in the XBB spike expression vector by QuickChange mutagenesis. The nucleotide sequence of all purified spike expression plasmids was verified by Sanger sequencing.

### HDX-MS Epitope Mapping

SARS-CoV-2 S-2P trimer at 3.33 μM was incubated overnight at 37 °C. The protein was then diluted to 1.66 μM trimer in the presence or absence of 6.25 μM R3DC23 in 1x PBS (pH 7.4, Sigma-Aldrich P4417). To initiate exchange, the protein was diluted tenfold into temperature-equilibrated deuterated buffer made by lyophilizing 1x PBS and resuspending in D_2_O (Sigma-Aldrich 151882). Samples were quenched at each time point (15s, 3m, 30m, 3h) by mixing 60 µl of the exchange reaction with 60 µl of ice-cold 2x quench buffer (3.6M guanidinium chloride, 500mM TCEP, 200mM glycine pH 2.4). The quenched samples were incubated on ice for 1 minute and then flash frozen in liquid nitrogen and stored at -80 °C until LC-MS. LC-MS and data analysis was conducted as previously described (40).

### SARS-CoV-2 challenge model in Syrian golden hamsters

In brief, 9-to 10-weeks-old male Syrian golden hamsters (Mesocricetus auratus) weighing 89.8 g to 132.3 g were obtained from Janvier (France). Housing conditions and experimental procedures were approved by the ethics committee in the Netherlands (study was registered under number 27700202114492-WP49). Six hamsters per group were infected intranasally with 100 TCID_50_/dose SARS-CoV-2 (Wuhan strain) in a total dose volume of 100 µl, divided equally over both nostrils. The test item huR3DC23-Fc_LS (2 and 10 mg/kg), palivizumab (10 mg/kg) and bebtelovimab (10 mg/kg) were administered by intraperitoneal injection 4 hours after the SARS-CoV-2 challenge. The irrelevant antibody palivizumab (Synagis, anti-RSV antibody) was used as a negative control, while bebtelovimab was used as a positive control. The huR3DC23-Fc_LS used in the hamster study was produced from stable cells pools. Hamsters were monitored daily for behavior, appearance and body weight.

On day 4 post-infection, animals were euthanized. At the time of necropsy, gross pathology was performed and abnormalities were recorded. Samples from the right lung lobes were collected and frozen for virological analysis. To determine virus titers, quadruplicate 10-fold serial dilutions were used in confluent layers of Vero E6 cells. To this end, serial dilutions of the samples (lung tissue homogenates) were incubated on Vero E6 monolayers for 1 hour at 37°C. Vero E6 monolayers were then washed and incubated for 5 or 6 days at 37°C, after which plates were stained and scored based on cytopathic effect (CPE) by using the vitality marker WST8 (colorimetric readout). Viral titers (Log_10_ TCID_50_/g) were calculated using the Spearman-Karber method. For the viral titration, the lower limit of detection (LLOD) ranged between 1.1 and 1.3 log_10_ TCID_50_/g. To detect viral RNA, lung tissue and homogenates were used. RNA was isolated and Taqman PCR was performed. The number of copies (Log_10_ CP/g) in the different samples was calculated against a standard included in each run. For the viral RNA, the LLOD was 3.5 Log_10_ CP/g.

Blood samples were collected prior to the start of the study on day -2 (∼ 200 μl of blood was collected for serum preparation under isoflurane anesthesia) and on day 4 post-infection (p.i.), at time of necropsy for pharmacokinetic analysis. Blood samples for serum preparation were immediately transferred to appropriate tubes containing a clot activator. Serum was collected and stored frozen. To inactivate potential infectious material present and to allow the testing of the sera samples in a BSL-2 environment, day 4 post-infection sera was heat-treated at 56 °C for 30 minutes.

### ELISA for detection of huR3DC23-Fc_LS in hamster sera

Pharmacokinetic analysis (PK) of the hamster serum samples was done using an ELISA based assay. Streptavidin-coated 96-wells microtiter plates were pre-blocked with superblock T20 (Thermo Scientific, catalogue number 37516) at room temperature. Afterwards, the plates were washed 3 times with 200µL of washing buffer (PBS with 0.05% Tween20). Biotinylated anti-VHH monoRab monoclonal antibody (Genscript, catalogue number A01995-200) was captured at a concentration of 0.1 μg/mL (10 ng/well in 0.1 mL) onto the streptavidin plate for 2 hours at room temperature with gentle shaking (400 rpm). Next, the supernatant was removed, and plates were washed three times with washing buffer. As a next step, 100 µl of 1.7-fold dilution series of standard calibrator (huR3DC23-Fc_LS), freshly prepared quality control samples (QCs) or hamster serum samples were added to the plates (according to a minimal required dilution of 10 or 50), which allowed huR3DC23-Fc_LS to bind to the captured anti-VHH antibody. After 1 hour incubation with shaking at room temperature, plates were again washed three times with washing buffer. To detect bound huR3DC23-Fc_LS onto the plates, anti-VHH monoRab antibody conjugated to horse-radish peroxidase (Genscript, catalogue number A01861-200) was added at a concentration of 0.2 µg/mL (0.1 mL per well) and incubated for 30 minutes at room temperature with shaking. Plates were washed three times with PBS buffer to remove the remaining Tween20. The final step was the addition of 100 µL of 1-Step Ultra TMB-ELISA Substrate solution (Thermo Scientific, catalogue number 34029) into wells for 5 minutes to allow a colorimetric reaction. Following the color development, the reaction was stopped by adding 50 µL of Stop solution (Thermo Scientific, Catalogue number N600). The colorimetric output was read on a Tecan Spark instrument with SparkControl software.

---

**Supplementary figure 1: Classification of neutralizing VHHs in subfamilies**. The amino acid sequences of neutralizing VHHs with a unique sequence were aligned, which was used to create an alignment tree. The dotted line represents the arbitrary cut-off to define subfamilies. All sequence analysis was done in CLC Main Workbench 22.0.1.

**Supplementary figure 2: S2 targeting VHHs potently neutralize SARS-CoV-2 pseudotyped virus on VeroE6/TMPRSS2 cells**. VeroE6/TMPRSS2 cells were infected with GFP reporter virus pseudotyped with SARS-CoV-2 VSV-S (D614G) that had been pre-incubated with a dilution series of the indicated VHH. VHH72-S56A was included as a positive and GFP-binding protein (GBP) as a negative control. The graphs show the mean ± SEM (N = 3) GFP intensity for each VHH dilution.

**Supplementary figure 3: S2 targeting VHHs potently neutralize replication-competent SARS-CoV-2 pseudotyped virus**. Vero E6 (left panel) or VeroE6/TMPRSS2 (right panel) cells were infected with replication-competent GFP reporter virus pseudotyped with SARS-CoV-2 VSV-S that had been pre incubated with a dilution series of the indicated VHH. VHH72-S56A was included as a positive and GFP-binding protein (GBP) as a negative control. The graphs show the mean ± SEM (N = 2) GFP intensity for each VHH dilution.

**Supplementary figure 4: Structural view of paratope – epitope contacts in the R3DC23 – HR2 complex**. **(A, B)** Close-up view of a single R3DC23 copy bound to two HR2 helices, with key interacting residues in the paratope and epitope shown in stick representation, colored as in Fig. 5. Candidate H-bonds and salt bridges are shown as red dashed lines. Escape mutant positions are labeled in red.

**Supplementary figure 5: Binding of huR3DC23-Fc_LS to FcRn at pH 6.0 and pH7.4**. **(A)** huR3DC23-Fc_LS showed improved steady state affinity to FcRn at pH 6.0 compared to IgG1K isotype bebtelovimab biosimilar control antibodies. Sensorgrams and steady state affinity plots for 7.8 – 250 nM of the indicated antibodies or 0.97 – 250 nM VHH-Fc binding to immobilized FcRn at pH 6.0. **(B)** Sensorgrams and steady state affinity plots for 7.8 – 1500 nM of the indicated antibodies or VHH-Fc show little binding to immobilized FcRn at pH 7.4. K_D_ was not determined as no steady state was reached even at 1500 nM antibody or VHH-Fc. **(C)** Steady state affinity and 1:1 binding model kinetic parameters. Conc: concentration of antibody or VHH-Fc; N/A: not applicable as steady state was not reached; *RU at maximum concentration of 1500 nM, no equilibrium.

**Supplementary figure S6: Detection of huR3DC23-Fc_LS in serum samples of challenged Syrian Golden hamsters**. Serum samples of hamsters that were treated with 10 mg/kg (A) or 2 mg/kg huR3DC23-Fc_LS (B) were collected at 4 days post challenge and tested by ELISA for the presence of huR3DC23-Fc_LS. As reference huR3DC23-Fc_LS was used and as negative controls serum of two hamsters treated with palivizumab were used. The graphs show mean ± SD (N = 2) OD_450_ (corrected with OD_650_) of 1.7-fold dilution series. The curves were fitted using non-linear fitting, [Agonist] vs. response -- Variable slope (four parameters). In the sera of animals 221186010150* and 221186010147* no or very low levels of huR3DC23-Fc_LS was detected.

**Table S1.** **Data collection statistics and refinement parameters for R3DC23-HR2**

|  |  |
| --- | --- |
| **Dataset** | **R3DC23-HR2** |
| **Data collection** |  |
| Synchrotron | Diamond Light Source |
| Beamline | i24 |
| Wavelength, Å | 0.61990 |
| **Data processing** |  |
| Space group | I222 |
| Cell parameters, Å<br>($\alpha=\beta=\gamma=90^\circ$) | 90.413 98.8 150.763 |
| Resolution, Å (outer shell)<br>(c) | 82.636 - 1.937<br>(2.052 - 1.937) |
| Total reflections | 543617 (26260) |
| No. of unique reflections | 39377 (1969) |
| Completeness (spherical) | 78.1 (24.9) |
| Completeness (ellipsoidal) | 95.2 (64.6) |
| Multiplicity | 13.8 (13.3) |
| $R_{\text{merge}}$, % | 0.274 (2.04) |
| $R_{\text{pim}}$, % | 0.076 (0.586) |
| $\langle I/\sigma(I) \rangle$ | 8.3 (1.5) |
| CC(1/2) | 0.990 (0.611) |
| Mosaicity, ° |  |
| **Refinement** |  |
| Resolution range, Å | 82.64 – 1.94 |
| No. of reflections | 39359 |
| Percentage observed | 78.02 |
| $R_{\text{cryst}}$, <sup>(a)</sup> % | 20.81 |
| $R_{\text{free}}$, <sup>(b)</sup> % | 25.09 |
| **RMS** |  |
| Bonds, Å | 0.024 |
| Angles, ° | 3.119 |
| **Ramachandran Plot** |  |
| Most favored, % | 95.56 |
| Additionally allowed, % | 2.22 |
| Disallowed, % | 2.22 |
| **PDB code** | XXXX |

<sup>[a]</sup> $R_{\text{cryst}} = S ||F_{\text{obs}}| - |F_{\text{calc}}|| / S |F_{\text{obs}}|$ , $F_{\text{obs}}$ and $F_{\text{calc}}$ are observed and calculated structure factor amplitudes
<sup>[b]</sup> $R_{\text{free}}$ as for $R_{\text{cryst}}$ using a random subset of the data excluded from the refinement
<sup>(c)</sup> Data in brackets are for the highest resolution shell

**Table S2:** **Neutralization of VSV pseudotyped with spike proteins of SARS-CoV-2 variants D614G, BA.4.6, BF.7, BQ.1.1, XBB, or XBB.1.5(-G252V) by huR3DC23-Fc_LS, sotrovimab (without LS) and bebtelovimab.**

| IC<sub>50</sub> ± SD (ng/mL) | huR3DC23-Fc_LS | sotrovimab (without LS)* | bebtelovimab* |
| --- | --- | --- | --- |
| D614G (n=5) | 0,8 ± 0,07 | 24,0 ± 1,27 | 1,4 ± 0,15 |
| BA.4.6 (n=3) | 0,6 ± 0,05 | 1076,0 ± 109,27 | 0,8 ± 0,07 |
| BF.7 (n=3) | 0,6 ± 0,08 | 1145,0 ± 167,59 | 0,8 ± 0,07 |
| BQ.1.1 (n=3) | 0,7 ± 0,07 | 1566,0 ± 232,41 | ND* |
| XBB (n=2) | 0,9 ± 0,14 | 282,0 ± 40,44 | ND* |
| XBB.1.5(-G252V) (n=3) | 0,5 ± 0,07 | 247,6 ± 24,30 | ND* |

The table shows the mean IC<sub>50</sub> values (ng/mL) that were calculated by nonlinear regression curve fitting, log(inhibitor) versus normalized response (four parameters). (\*) bebtelovimab biosimilar and sotrovimab (without LS) biosimilar were used as positive controls. (ND\*) No IC<sub>50</sub> value could be calculated due to absence of neutralization activity within the tested concentration range (0.0064 – 20 ng/ml) of bebtelovimab biosimilar. The viruses for which the tested antibodies have lost <5-fold, 5<25-fold and >25-fold of neutralization activity are respectively indicated in green, yellow and red.

## Acknowledgements

We are grateful to G. Zimmer providing reagents to generate VSV pseudotype particles; J. McLellan and D. Wrapp for providing SARS-CoV-2 spike and RBD proteins and expression plasmids for these proteins; We thank L. de Waal from Viroclinics for coordinating the hamster challenge study. We thank Piet Maes for providing a SARS-CoV-2 Omicron BA.1 isolate and Florian I Schmidt for providing replication competent VSV pseudotyped with SARS-CoV-2 spike. We thank the staff of the VIB Flow Core Ghent for providing access to flow cytometry equipment and the support with flow cytometry experiments. We also thank the staff of the VIB Bioimaging Core Ghent for outstanding support with microscopy imaging. We acknowledge all Global Initiative on Sharing All Influenza Data (GISAID) contributors for sharing the sequencing data. S.M. is a Chan Zuckerberg Biohub Investigator. We thank the National Institute for Biological Standards and Control (NIBSC) for providing VeroE6/TMPRSS2 cells. This work was in part executed by the ad hoc VIB Center for Medical Biotechnology COVID-19 emergency drug development team.

