## Supplementary figures for "Ultrapotent SARS coronavirus-neutralizing single-domain antibodies that bind a conserved membrane proximal epitope of the spike"

Supplementary figure 1: phylogenetic tree of R3DC23 related VHHs

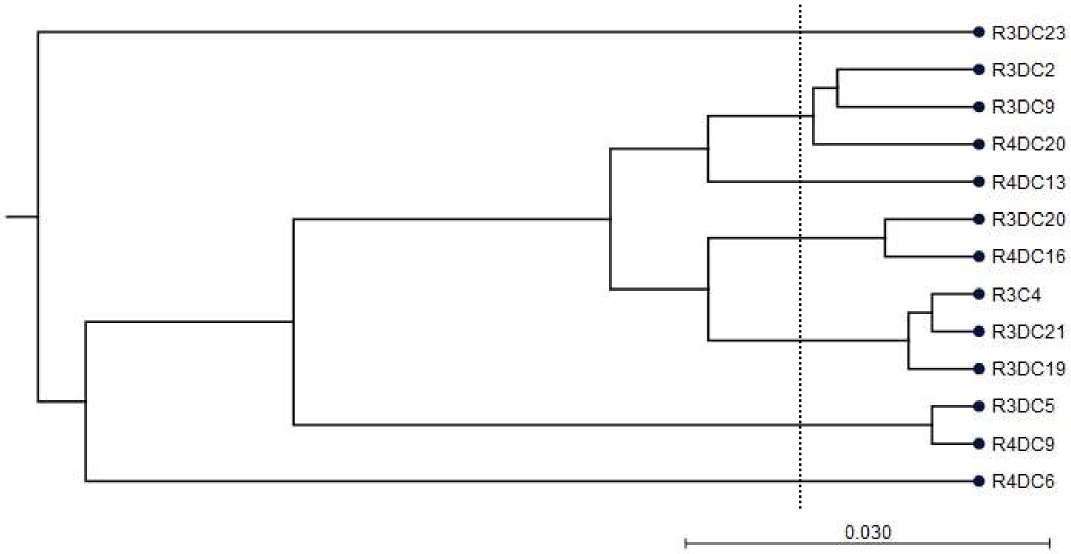

### Supplementary figure 2:neutralization on VeroE6/TMPRSS2 cells

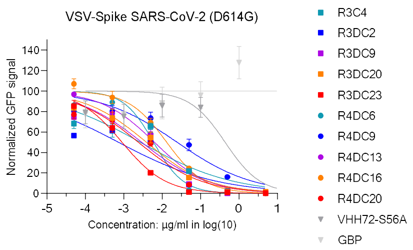

Supplementary figure 3:
Neutralization of replicating VSV-Spike on Vero E6 and VeroE6/TMPRSS2 cells

A

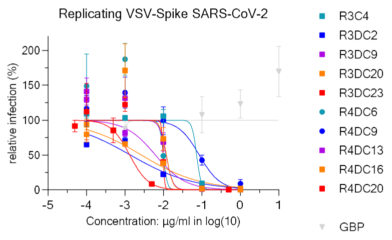


B

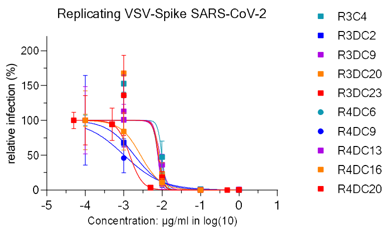

Supplementary figure 4:
Structural details on the R3DC23/HR2 interaction

A

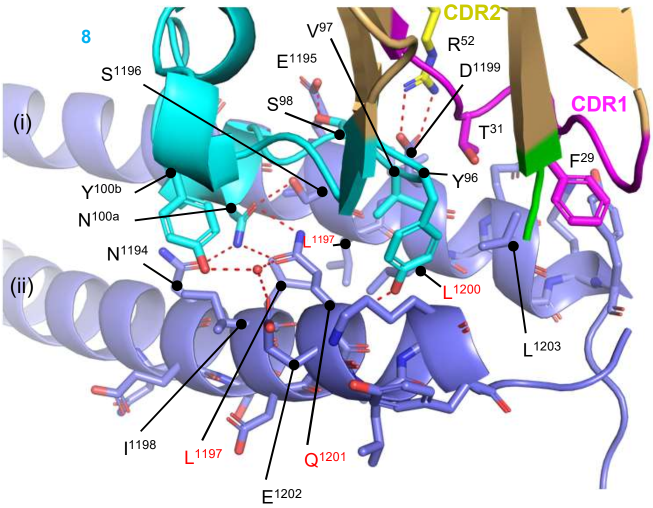


B

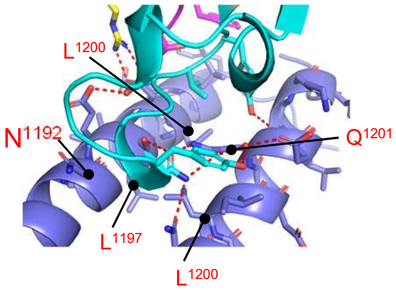

### Supplementary figure 5:
### Binding of huR3DC23-Fc_LS to FcRn at pH 6.0 and pH 7.4

#### A Binding of antibodies to FcRn at pH 6.0

bebtelovimab biosimilar | hIgG1K isotype control | huR3DC23-Fc_LS

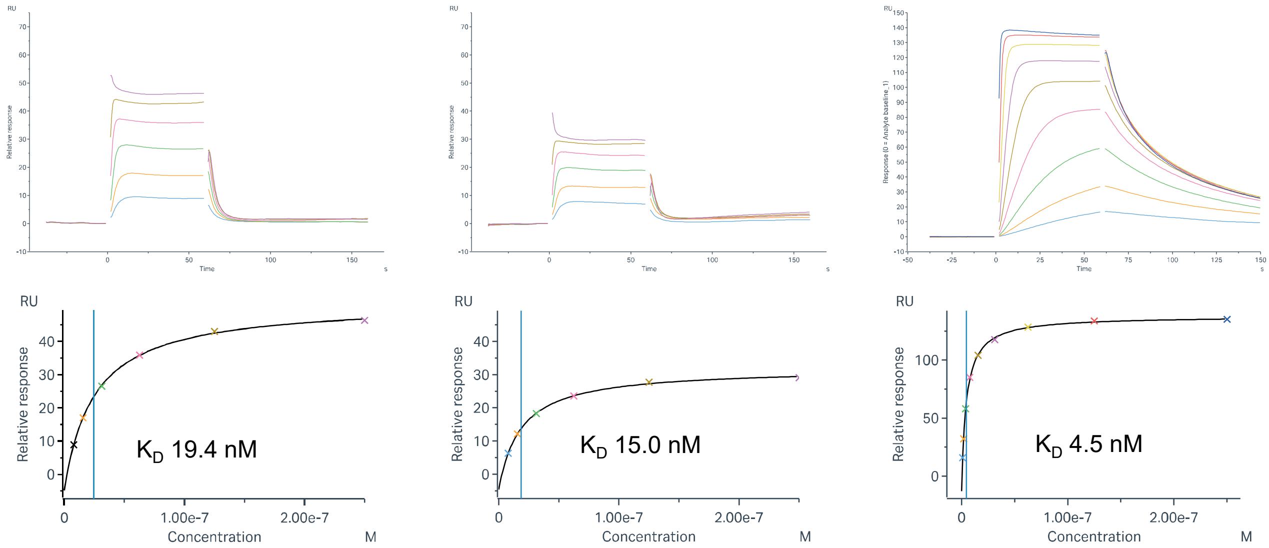


#### B Binding of antibodies to FcRn at pH 7.4

bebtelovimab biosimilar | hIgG1K isotype control | huR3DC23-Fc_LS

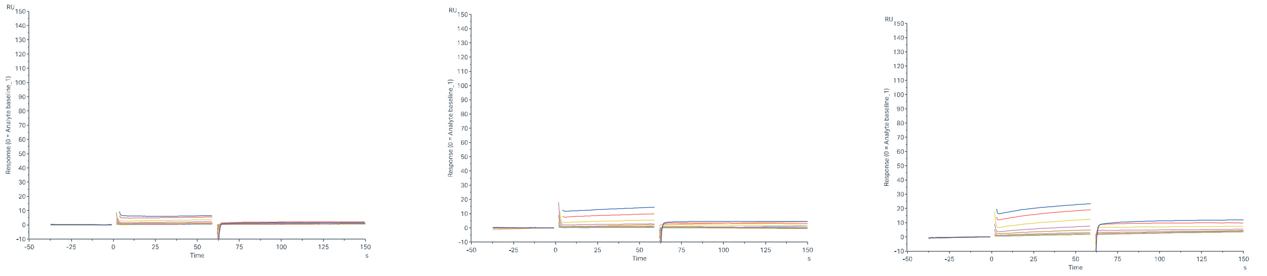

#### C Binding characteristics of antibodies to FcRn at pH 6.0 and pH 7.4

| | FcRn pH 6.0 | | | | | FcRn pH 7.4 | |
|---|---|---|---|---|---|---|---|
| | | Steady state | | 1:1 binding model | | | |
| | Conc (nM) | $RU_{max}$ (RU) | $K_D$ (nM) | $RU_{max}$ (RU) | $K_D$ (nM) | Conc (nM) | $RU_{max}$ (RU) |
| huR3DC23-Fc_LS | 250 – 0.97 | 149.8 | 4.46 | 135.2 | 3.5 | 1500 – 7.8 | 22.4* |
| Bebtelovimab biosimilar | 250 – 7.8 | 56.6 | 24.7 | N/A | N/A | 1500 – 7.8 | 8.2* |
| hIgG1K isotype control | 250 – 7.8 | 36.5 | 18.5 | N/A | N/A | 1500 – 7.8 | 16.2* |

### Supplementary figure 6:
### Detection of huR3DC23-Fc_LS in the serum of hamsters treated With 10 or 2 mg/kg huR3DC23-Fc_LS

A

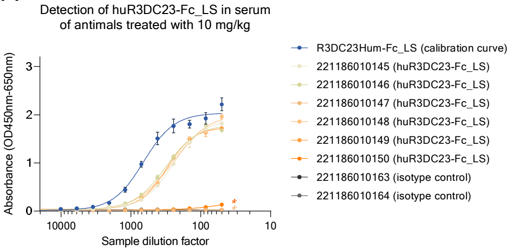


B

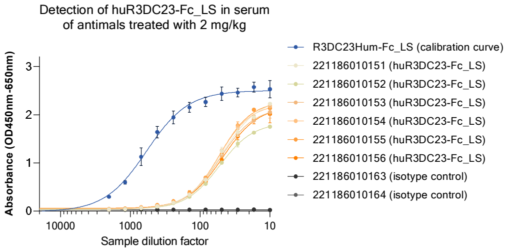
